# A comparison of lipid diffusive dynamics in monolayers and bilayers in the context of interleaflet coupling

**DOI:** 10.1101/2024.04.26.589162

**Authors:** Titas Mandal, Nadine Brandt, Carmelo Tempra, Matti Javanainen, Balázs Fábián, Salvatore Chiantia

## Abstract

Cellular membranes are composed of lipids typically organized in a double-leaflet structure. Interactions between these two leaflets – often referred to as interleaflet coupling – play a crucial role in various cellular processes. Despite extensive study, the mechanisms governing such interactions remain incompletely understood. Here, we investigate the effects of interleaflet coupling from a specific point of view, i.e. by comparing diffusive dynamics in bilayers and monolayers, focusing on potential lipid-specific interactions between opposing leaflets. Through quantitative fluorescence microscopy techniques, we characterize lipid diffusion and mean molecular area in monolayers and bilayers composed of different lipids. Our results suggest that the observed decrease in bilayer lipid diffusion compared to monolayers depends on lipid identity. Furthermore, our analysis suggests that lipid acyl chain structure and spatial configuration at the bilayer may strongly influence interleaflet interactions and dynamics in bilayers. These findings provide insights into the role of lipid structure in mediating interleaflet coupling and underscore the need for further experimental investigations to elucidate the underlying mechanisms.

## 1 Introduction

Cell membranes consist of bilayers ^1^ whose structural integrity and function depend on the self-assembly of a plethora of structurally diverse lipid molecules ^2,3^. Furthermore, cellular lipid bilayers possess nano- to microscale heterogeneities (i.e., domains) with different characteristics than the surrounding bulk membrane ^4–6^. Fluorescence microscopy studies of nano- (< ca. 200 nm) and microdomains (> ca. 200 nm) in model membranes have shown that these entities do not exist just in one leaflet, but rather span the whole bilayer ^4,7–9^. In this context, the interaction between the two leaflets in lipid membranes (i.e., interleaflet coupling ^10–12)^ has been shown to be important for various cellular processes ^13^, including e.g. signal transduction ^14–18^, registered lipid-protein domain formation ^12,19^ and host-pathogen interactions ^14,15,20–22^. A common feature of these phenomena is that lipid-lipid interactions (e.g., packing, phase separation, diffusive dynamics) are somehow communicated to or shared between the opposing leaflets in a bilayer.

Interleaflet coupling has been experimentally characterized from different points of view and within different systems ^13^. For example, several groups have investigated how lipid domain formation in one leaflet can induce analogous phase separation in the opposing leaflet ^23–26^ or, more in general, how phase transitions are coupled between leaflets ^27^. Alternatively, the friction or the shear stress between the two leaflets have been analysed both theoretically ^28–30^ and experimentally ^31–33^. Finally, several efforts have been made to understand how lipid dynamics in one leaflet affect those in the other leaflet, including investigations via fluorescence spectroscopy ^34–37^. In this context, we and others have observed that the interleaflet coupling of lipid diffusive dynamics in symmetric and asymmetric phosphatidylcholine bilayers (e.g. OMPC, POPC and SOPC) might depend on their acyl chain structure ^34,38^.

Several physical processes have been proposed to mediate interleaflet interactions, including membrane undulations, cholesterol flip-flop, line tension and interdigitation (see the review by Sarmento *et al.* ^13^ and references within). While the latter mechanism is controversially discussed in the literature ^10,39,40^, it is reasonable to hypothesize that interleaflet coupling might be indeed mediated by interactions at the bilayer midplane between acyl chains belonging to opposing leaflets ^13,34,38,40,41^.

Here, we focus on a specific possible manifestation of interleaflet coupling, i.e. the decreased lipid diffusive dynamics observed in bilayers, compared to monolayers. Lipid monolayers (e.g., at air-water ^42^ interfaces) are simple biophysical models of each leaflet in a lipid bilayer and are characterized in general by fast lipid diffusion (25-35 µm²/s) ^43–45^. Free-standing lipid bilayers, on the other hand, exhibit lateral diffusion coefficients (*D*) for lipids with values between ca. 5 and 12 µm²/s ^34,46,47^. In the context of the Saffman-between lipid monolayers and bilayers. Any further decrease of lipid diffusion in bilayers might be attributed to some form of interleaflet interactions ^42,45^. In this work, we have performed a systematic analysis of this effect by comparing monolayers and bilayers composed of different lipids. Lipid diffusion and mean molecular area (*MMA*) in monolayers and bilayers were quantified using fluorescence fluctuation microscopy methods (i.e. line-scan fluorescence correlation spectroscopy (lsFCS) ^49^ and raster image correlation spectroscopy (RICS) ^50^. Both these approaches have been previously used to characterize the physical properties of model membranes, such as giant unilamellar vesicles (GUV) ^34,47^ and lipid monolayers ^43,51^.

By analysing the behaviour of lipids which were previously shown to potentially influence interleaflet coupling ^34,38^, we aimed to verify the hypothesis that diffusive dynamics in lipid bilayers might be determined by lipid-specific interactions between opposing leaflets.

## 2 Materials and Methods

### 2.1 Chemicals

For the preparation of lipid monolayers and GUVs, the following lipids were purchased from Avanti Polar Lipids (Alabaster, AL, USA): 1-oleoyl-2-myristoyl-sn-glycero-3-phosphocholine (OMPC), 1,2-dioleoyl-sn-glycero-3-phosphocholine (DOPC), 1-palmitoyl-2-oleoyl-glycero-3-phosphocholine (POPC), 1-stearoyl-2-oleoyl-sn-glycero-3-phosphocholine (SOPC), Sphingomyelin (Milk, Bovine), 1-oleoyl-2-(6-((4,4-difluoro-1,3-dimethyl-5-(4-methoxyphenyl)-4-bora-3a,4a-diaza-s-indacene-2-propionyl)amino)hexanoyl)-sn-glycero-3-phosphocholine (TF-PC) and 1,2-distearoyl-sn-glycero-3-phosphoethanolamine-N-(lissamine rhodamine B sulfonyl-ammonium salt/ Rh-PE). 10X Phosphate Buffer Saline (pH 7.4) for GUV experiments, Glycerine trioleate (Triolein) for the preparation of lipid droplets, Alexa Fluor^®^ 488 dye and Alexa Fluor^®^ 555 dye for the calibration of excitation beam waist, isopropanol for monolayer glass chamber cleaning were purchased from ThermoFisher Scientific (Waltham, MA, USA). Sucrose for electroformation was purchased from PanReac Applichem GmbH (Germany). Bovine Serum Albumin (BSA) for GUV chamber coating, Disodium phosphate and Monosodium phosphate for Sodium phosphate buffer (NaP) preparation and ethanol for monolayer glass chamber cleaning were purchased from Carl Roth GmbH (Germany). Hellmanex^®^ for monolayer glass chamber cleaning was purchased from Hellma GmbH & Co. KG (Germany).

### 2.2 Lipid mixing and air-water monolayer preparation

Lipid solutions were mixed in glass vials using positive-displacement micropipettes with glass capillary tubes. Methanol was used as solvent and a minimum volume of ∼1 mL was used for all lipid solutions. These precautions mitigated concentration errors due to pipetting and solvent evaporation. Lipid solutions were stored in glass vials sealed with polytetrafluoroethylene (PTFE) tape and wrapped in aluminium foil.

Lipid monolayers were prepared similarly to what was described before ^42,43^. To this aim, lipid solutions were always prepared at 0.1 mg/mL and contained 0.005 mol% TF-PC. This dye concentration was optimized for best signal to noise (S/N) ratio in fluorescence correlation measurements. The effective concentration of TF-PC was confirmed after each lipid mixture preparation via spectrophotometry measurements of serial dilutions. Then, the solvent was dried with a nitrogen stream and replaced with chloroform. This facilitates the solvent evaporation step needed after spreading the lipid solution at the air-water interface. Prior to each experiment, the monolayer chamber ^43^ was cleaned using sequentially 2% Hellmanex^®^ in water, isopropanol, ethanol, and Milli-Q^®^ water. The cleaned chamber was dried with compressed air after every rinsing step. To prepare lipid monolayers, 200 µl of autoclaved and filtered Milli-Q^®^ water was added to the chamber and the lipid solution in chloroform was spread dropwise on the surface. The volume of the solution depended on the desired *MMA* (*MMA_expected_*) of the monolayer to be obtained, according to the following equation:

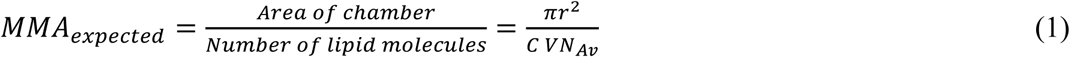

wherein *r* is the radius of the monolayer chamber, *C* is the concentration of the 0.1 mg/mL lipid solution in molar units, *V* is the volume to be spread and *N_Av_* is the Avogadro’s number. After solvent evaporation (∼15 mins), 30-50 µl of water was carefully removed to keep the monolayer within the working distance of the objective ^43^. Finally, the chamber was carefully covered with a Teflon coated lid to minimise water evaporation. All measurements were performed at room temperature.

### 2.3 GUV preparation

GUVs with different lipid compositions were prepared by electroformation ^52,53^ using cylindrical Teflon chambers containing two platinum wires ^52,53^. Lipid solutions (2.5 mg/mL with 0.005 mol% TF-PC) were prepared in ethanol and 2 µL were spread evenly on each wire twice and then dried under nitrogen stream. TF-PC concentration was optimized for best S/N ratio in lsFCS measurements. GUV swelling took place in the Teflon chamber filled with a 50 mM sucrose solution, applying alternating voltage (2 V peak-to-peak, 10 Hz) for 1 hour. To detach the GUVs from the wires, the process was continued for 30 minutes longer, decreasing the frequency to 2 Hz. The entire procedure was performed at room temperature. After electroformation, 100 µL of GUV suspension was transferred to custom-made microscopy chambers, pre-treated with a 10% BSA solution for 15 minutes. The GUV suspension was further diluted with an equal volume of 40 mOsmol/kg PBS containing 0.3% (w/v) low melting agarose to induce a slight inflation and stabilization of GUVs ^46^.

### 2.4 Lipid droplet preparation (oil-water monolayers)

Lipid droplets (LDs) were prepared with Triolein as oil-phase according to available protocols ^54,55^. A lipid solution doped with 0.001 mol% Rh-PE in methanol was dried in microcentrifuge tubes under a nitrogen stream. Unlike the case of GUVs and air-water monolayers, TF-PC showed a significantly low fluorescence signal in oil-water monolayers if similar excitation powers were used (data not shown). Moreover, higher excitation intensities resulted in extensive photobleaching. For these reasons, the fluorescent head-labelled lipid Rh-PE was used as a probe for experiments in LDs, rather than TF-PC. Pre-warmed triolein was added to the dried film (500:1 mass ratio) and the mixture was extensively vortexed for 15 minutes and sonicated for 30 minutes. The solution was then added to 20 mM NaP buffer of pH 7.4 (1:10 v/v) and sonicated for 10 seconds after short vortexing, thus obtaining an oil in buffer emulsion that contained the LDs. 75 µL of this LD emulsion was transferred to a single well of a 96-well plate and further diluted with an equal volume of the same NaP buffer prior to imaging. This procedure results in LDs with low mobility (as they appear to be adhering to the glass surface of the observation chamber).

### 2.5 Line-Scan Fluorescence Correlation Spectroscopy

Line-Scan Fluorescence Correlation Spectroscopy (lsFCS) was performed on GUVs and LDs as previously described ^49,56–58^. Briefly, GUVs or LDs were scanned perpendicularly 500,000 times using a pixel dwell time of 0.79 µs and resolution of 256 pixels (total scan time ∼4 min). The pixel size was 0.08 µm. A 488 nm argon laser at a power of 1.45 µW was used to excite the GUV samples and a 561 nm laser (6.5 µW) was used to excite the LD samples, while keeping the photobleaching below ca. 10% of the total initial intensity. Each GUV or LD was scanned only once. All measurements for the GUVs were performed on a Zeiss LSM780 system (Carl Zeiss, Oberkochen, Germany) using a Plan-Apochromat 40x/1.1 Korr DIC M27 water immersion objective. Measurements on the LDs were performed on a Zeiss LSM880 system with Plan-Apochromat 40x/1.2 Korr DIC M27 water immersion objective. Calibration of the excitation beam waist (*w_0_*) was performed each day by measuring the fluorescence autocorrelation curve of 0.25 µM Alexa Fluor^®^ 488 dye in 40 mOsmol/kg PBS with 35 mM sucrose for the GUV experiments to correct the variation in laser alignment. The same calibration procedure was performed for LD experiments, using a similar concentration of Alexa Fluor^®^ 555 dye in 20 mM NaP buffer. A mean of the diffusion times *(τ_D_*) from three independent autocorrelation curves (100 repetitions with acquisition time 5 seconds) was used to calculate *w_0_* using the reported diffusion coefficient (*D*) of the fluorescent dyes ^59,60^ neglecting, as an approximation, the ca. 2% decrease due to the presence of the solute ^61,62^:

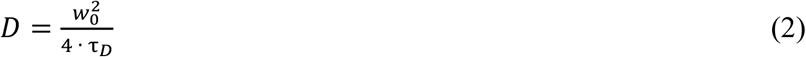

Additionally, the same calibration procedure was used to verify that the number of detected particles (*N*) in the confocal volume (*π^3/2^· w_0_^2^·z_0_*, in which *w_0_* and *z_0_* are the axial and radial sizes of the confocal volume) matched the known dye concentration ^63^.

All lsFCS measurements were performed at room temperature and with a confocal pinhole size of 39 µm for measurements in GUVs and 45 µm for those in LDs. The collar ring of the objective was adjusted to optimize the signal. Each data set was exported as TIFF files and then analysed in MATLAB (The MathWorks, Natick, MA) using custom-written code ^64^.

The effective illuminated area (*A_eff_*) measured on the GUV or LD during the line scanning was calculated as^49^:

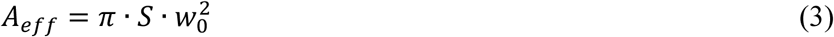

wherein *S* is the structural parameter. *S* was estimated using two approaches. First, we tried to determine this value as fit parameter of the calibration autocorrelation curve measured for the free dye in solution (see above). This approach is characterized in general by poor reproducibility (data not shown). Therefore, as an alternative, we measured autocorrelation curves at the north pole of a limited set of GUVs for each lipid composition using point FCS ^51^, for which the *A_eff_* is independent of *S* (*A*_*eff*_ = *π* · *w*^2^) ^65^. The same GUVs were also measured via lsFCS at the equatorial plane. Given that *N* is in both cases equal to *A_eff_* times the fluorescent lipid concentration, the ratio of *N* obtained from lsFCS to that obtained from point FCS yielded *S*. Median values obtained from the two approaches were comparable (∼7) and *S* was therefore fixed to 7.

Each lsFCS measurement provided the number of labelled lipids *N* in the measured area (calculated simply as the inverse of the autocorrelation amplitude, neglecting the ∼1% background signal) and *τ_D_* for TF-PC or Rh-PE. A simple calculation yielded the actual number of all (i.e., labelled and unlabelled) lipid molecules in the illuminated area (*N_total_*), since the dye was included at a known molar concentration compared to the other lipids. Finally, the mean molecular area (*MMA_b_*) or area per lipid in the bilayer was calculated using equation 4:

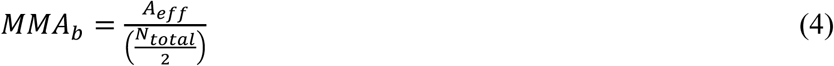

In LDs, the mean molecular area (*MMA_m-oil_)* was calculated as:

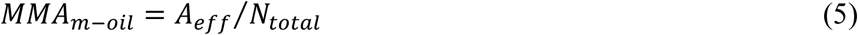

### 2.6 Raster Image Correlation Spectroscopy

Raster Image Correlation Spectroscopy (RICS) was performed on air-water lipid monolayers as previously reported ^50,66^. The microscope setup was the same as it was for the GUV measurements described in the previous section. A total of 30 frames with a resolution of 256 x 256 pixels (pixel size 0.05 µm) were acquired at a pixel dwell time of 1 µs and 2.2 µW laser excitation power. The photobleaching was thus always below 10% of the initial intensity. Due to continuous evaporation of the water subphase, the monolayer moved counteract this, at the beginning of the measurement, the focus was positioned ca. 1 µm below the monolayer to allow the focal plane to move in the opposite *z*-direction and the measurement time was kept short (ca. 4 seconds) to get a quasi-constant and maximized signal ^67^. Each data set was analysed in MATLAB using custom-written code ^68^. The analysis directly provided lipid diffusion in monolayers (*D_m_*) and the surface concentration *C* of TF-PC molecules in the monolayer. For the analysis, the background signal was neglected as it was generally less than 1% of the main signal. As described in the previous section (section 2.5), knowledge of TF-PC concentration relative to non-fluorescent lipids (*C_%TF-PC_*) and *C* allowed the estimation of the lipid *MMA* (*MMA_m_*):

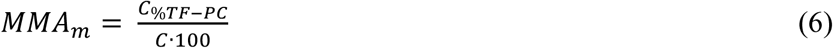

The correct formation of a lipid monolayer with the expected properties was also confirmed at this step by comparing *MMA_m_* with the anticipated value of *MMA_expected_* obtained from equation 1. Similar *MMA_m_* and *MMA_expected_* values for each monolayer sample indicated proper segregation of all lipids at the air-water interface.

### 2.7 Molecular dynamics simulations

To calculate the thicknesses of the monolayers and bilayers and to study the effect of acyl chain chemistry on interleaflet interactions, we resorted to all-atom molecular dynamics (MD) simulations. Single-component lipid bilayers containing a total of 1024 POPCs, DOPCs, SOPCs, or OMPCs and solvated by 50 water molecules per lipid were set up using CHARMM-GUI and equilibrated following the associated protocol ^69^. While POPC, DOPC, and SOPC are readily available in the CHARMM lipid library, the OMPC topology was built in-house following the building block approach of CHARMM lipids. These bilayers were then simulated for 100 ns, after which their MMAs had stabilised. We then converted the bilayer systems into monolayer ones by the translation of the coordinates of one of the leaflets and the simultaneous addition of a vacuum slab into the system. The monolayer systems contained two monolayers lining the water slab, and a vacuum phase (∼17.5 nm) separating the acyl chains across the periodic boundary conditions.

The lipids were modelled using the CHARMM36 lipid force field ^70^. Water was described with the CHARMM-specific TIP3P potential ^71,72^. All four bilayers and four monolayers were simulated for 1 µs using GROMACS 2021.5 ^73^, and the last 900 ns were used in all analyses.

The integration of the equations of motion was performed with a leap-frog integrator with a time step of 2 fs. The smooth particle mesh Ewald algorithm was used to incorporate long-range electrostatics with the real space cutoff optimised for each run ^74^. The Lennard-Jones potential was truncated at 1.2 nm, and the forces were switched to zero beyond 1.0 nm. Information of atomic neighbours was maintained using buffered Verlet lists ^75^. The monolayers were simulated in the canonical (NVT) ensemble, whereas the isothermal–isobaric (NPT) ensemble was used for the bilayers. The temperature of the lipids and water were separately coupled to the Bussi–Donadio–Parrinello thermostat ^76^. The target temperature was set to 298.15 K, and the time constant to 1 ps. For bilayers, pressure was maintained at 1 bar using a semi-isotropic coupling scheme (the two dimensions in the plane of the membrane coupled together). The Parrinello–Rahman barostat ^77^ with a time constant of 5 ps and compressibility of 4.5×10^-5^ bar^-1^ was used. All bonds involving hydrogen atoms were constrained using P-LINCS ^78^.

In order to estimate monolayer and bilayer thickness values with a single approach, we considered the bilayer and monolayer thickness to be the span of the *z* coordinate (along the bilayer/monolayer normal) in which the lipid density was larger than 10% of its maximum value (see supplementary Figure S1). These thicknesses were evaluated for the bilayer and the monolayer, as well as for a single leaflet in the bilayer. The density profiles were extracted using the gmx density tool included in the GROMACS distribution.

### 2.8 Viscosity measurements and interleaflet coupling

The aim of this work is to compare interleaflet interactions for different lipids, by quantifying the difference in lateral diffusion between monolayers and bilayers. However, different lipid compositions result in varying membrane thickness values which, in turn, would affect diffusion dynamics ^48,79^. For this reason, we quantify the viscosity of lipid membranes and interleaflet interactions by evaluating the experimentally obtained *D* and thickness values in the context of the Saffman-Delbrück-Hughes-Pailthorpe-White (SDHPW) model ^48,79^. To this aim, we prepared air-water lipid monolayers and LDs (i.e., oil-water monolayers) with the same *MMA* of the corresponding bilayers (i.e. bilayers with the same lipid composition), as mentioned in section 2.3. Such *MMA* values are in the range 60-70 Å^2^/lipid (see below). The experimentally obtained lipid diffusion in monolayers (*D_m_*) and the monolayer thickness *h_m_* obtained from simulations were then used to calculate the three-dimensional monolayer bulk viscosity *µ_m_* according to the SDHPW model (see Table S1). Using the same formalism of Vaz *et al*. ^80^, the reduced radius is defined in this case as:

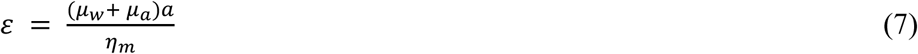

with

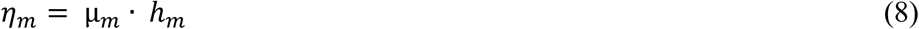

wherein *η_m_* is the monolayer surface viscosity, *a* is the radius of the diffusing lipid probe and the viscosities of the bounding fluids on both sides of the sheet i.e., *µ_w_* and *µ_a_* are those of water ^81^ and air ^82^, respectively. To avoid complications, we have assumed the radius *a* of the fluorescent lipid probe to be similar to the radius of the non-fluorescent lipid molecules ^83,84^. Since, for our experiments, the reduced radius *ε* is not much smaller than 1 in general, we make use of the analytical approximation introduced by Petrov *et al*. ^85^ to determine viscosity (*µ*) from *D*. Similar calculations are also performed for monolayers at the oil-water interface: from *D_m-oil,_* it is thus possible to estimate the bulk viscosity of the monolayer *µ_m-oil_*, considering that the viscosities of the bounding fluids on both sides of such monolayers i.e., *µ_w_* and *µ_t_* are those of water and triolein ^86^, respectively. Of note, only single lipid mixtures resulted in reproducible and stable LD preparations, and, for this reason, oil-water monolayers were prepared exclusively from either DOPC, POPC, SOPC or OMPC samples. For simplicity, we assume here that *h_m_* does not change considerably between the two types of monolayers.

Differently from the case of lipid monolayers, the application of SDHPW-related models to study the diffusion of lipids in bilayers (i.e., objects spanning only one of two leaflets) is less univocally described in the literature. On the one hand, the relationship between the thickness of the sheet (i.e., the bilayer membrane thickness, *h_b_*), the thickness of a leaflet *h_l_* and the length of the diffusing object is not clear ^28,87^. Additionally, although continuum models are used with great utility for modelling small solute diffusion ^28^, it was suggested that other models (e.g., free volume model ^80^) might describe lipid diffusion in bilayers better than a continuum fluid hydrodynamic formalism. Nevertheless, it must be noted that these models have several parameters that are difficult to independently quantify in the context of a comparison of lipids with different structures. Most importantly, the goal of this work is not to provide an absolute estimation of viscosity values, but rather to compare the decrease in lipid diffusivity between monolayers and bilayers, for different lipid types. Therefore, we compare here monolayers and their corresponding bilayers (i.e., same lipid composition and similar *MMA*). It is worth noting that the lateral tension differs strongly between these systems, as bilayers have vanishingly small tension ^88^ and monolayers require a lateral tension of ∼30 mN/m ^89^ in order to be comparable to bilayers. First, following the reasoning of Vaz *et al*. ^80^, TF-PC in a bilayer is assumed to be embedded and diffuse within a single leaflet with bulk viscosity *µ_l_*. The viscosities of the bounding fluids are assumed to be that of water, *µ_w_*, on one side of the leaflet and that experienced at the bilayer midplane, *µ_midplane_*, on the other. Using the experimentally determined values for lipid diffusion in bilayers (*D_b_*) and *h_l_* from MD simulations, we calculated *µ_midplane_*, using the analytical solution by Petrov *et al*. ^85^ and the following definition ^80^:

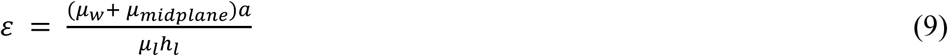

For each lipid composition, several replicate measurements on monolayers and bilayers were performed. For each estimation of *D_m_*, one *µ_m_* value was calculated. From each *µ_m_* value, one *µ_midplane_* value was calculated using a single *D_b_* value (i.e., <*D_b_*> calculated for the specific lipid composition). For example, as shown in Table S1, in the case of OMPC, 49 measurements of *D_m_* resulted in 49 *µ_m_* values, 49 *µ_midplane_* values and one *D_b_* value (obtained, in turn, as average of 122 measurements on bilayers). Vanishingly small *µ_midplane_* values (i.e., comparable to *µ_a_*) are obtained if the diffusivity in (a leaflet of) the bilayers was approaching that measured in the corresponding air-water monolayer, i.e. if each leaflet was “unaware” of the presence of the other. For this reason, we use *µ_midplane_* as an approximate measure of the interleaflet interaction, under the simplifying assumption that *µ_l_* is not affected by such interactions (i.e., *µ_l_* = *µ_m_*).

As an alternative, we followed one of the approaches mentioned by Adrien *et al.* ^87^. Briefly, in a rough approximation, lipid-like fluorescent probes can be treated as cross-layer particles, since it was often observed that *D* only weakly depends on the length of the diffusing lipid ^28,29,33,90–93^. This implies that the length of the diffusing molecule is simply assumed to be equal to the thickness of the whole bilayer *h_b_*. In the absence of any additional interaction between the leaflets, the bilayer bulk viscosity *µ_b_* should be the same as the bulk viscosity measured in the corresponding monolayer. The bulk viscosities of the bounding fluids are that of water, *µ_w_*, on one side and that of 50 mM sucrose, *µ_s_*, on the other side of the membrane:

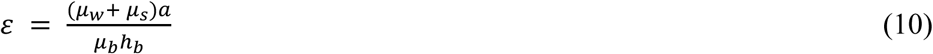

Using each experimentally derived *D_b_* values and *h_b_* (from lsFCS measurements and MD simulations, respectively), we calculated the bilayer bulk viscosity *µ_b_*, for each lipid composition. For simplicity, the bulk viscosity of 40 mOsmol/kg PBS is considered to be the same as that of water. If *µ_b_* were in fact equal to *µ_m_*, *D_b_* would be lower than *D_m_* simply due to the increased thickness of the lipid sheet and the presence of water on both its sides (instead of water and air). Any further decrease in *D_b_* (compared to *D_m_*) is taken here as an indication of coupling/ interaction between the two leaflets, as previously suggested ^42^. Concretely, this is estimated as the ratio <*µ_b_>*/*µ_m_* or <*µ_b_>*/*µ_m-oil_* (referred to as “coupling factor”). For example, as shown in Table S1, in the case of OMPC, 49 *µ_m_* values were experimentally obtained from measurements in monolayers and 49 values for the corresponding coupling factor were calculated, using a single <*µ_b_>* value (obtained as the average of 122 measurements on bilayers). Regarding the interpretation of the coupling factor, a value of ca. 1 would correspond to the above-mentioned case in which *D_b_*< *D_m_* only because of increased thickness and surrounding fluid viscosity. Concretely, for the lipid systems investigated here, this would correspond to a ca. 2-fold reduction in *D* when switching from air-water monolayers to bilayers. A coupling factor value above one would indicate that the leaflets in a bilayer have a higher bulk viscosity than their monolayer counterpart and we attribute this increase to interleaflet interaction.

### 2.9 Statistical analyses

Box plots in Figures 1 and 3 represent the mean, median, first and third quartile with standard deviations as whiskers. Bars in Figures 4, S3 and S4 represent mean values with standard deviations as whiskers. Each point in Figure 2 represents mean values with standard error of the mean as error bars. Statistical significance between different datasets for all figures was determined using two-sample t-test (significance level=0.05). All figures and statistical tests were made using Origin (Pro) version 2023 (OriginLab Corporation, Northampton, MA, USA).

**Figure 1:**
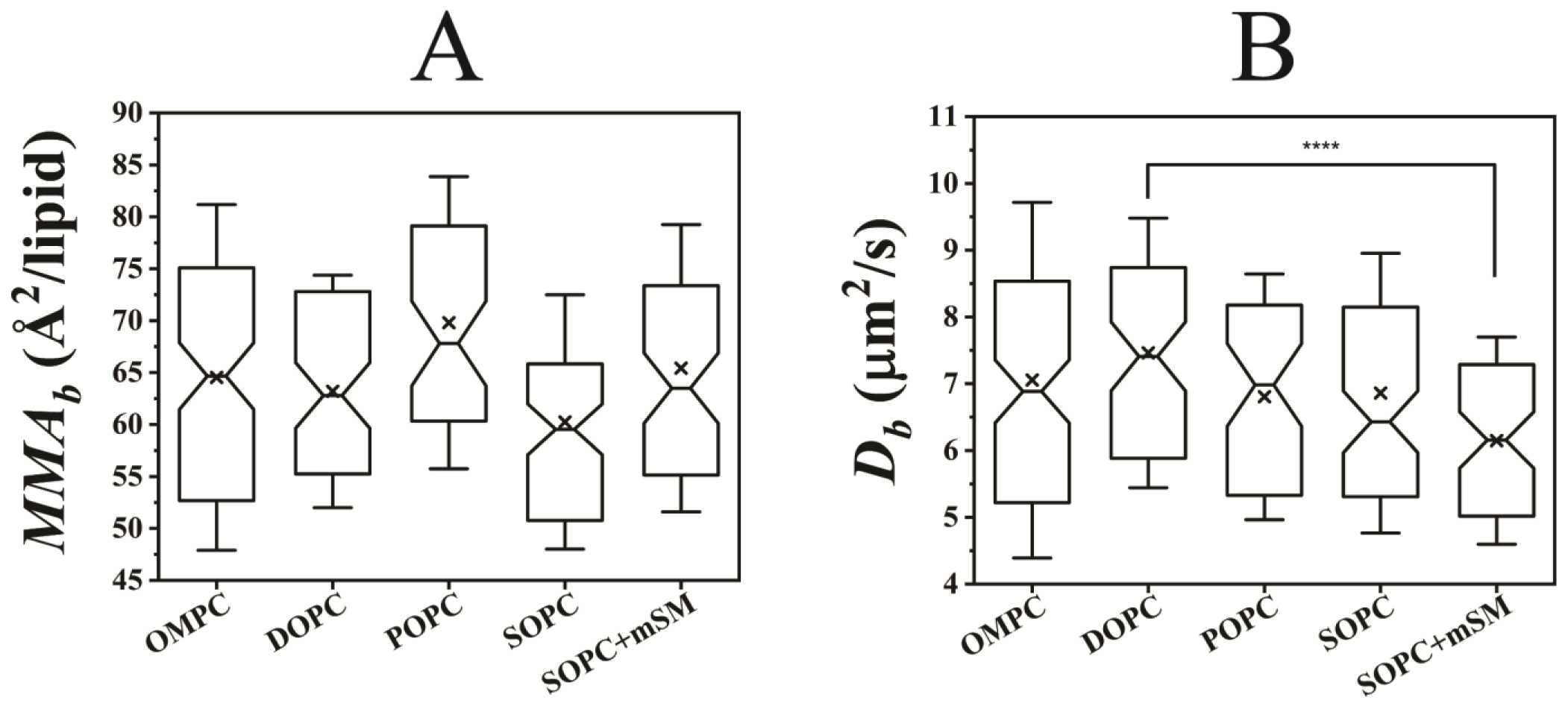
*MMA_b_* and *D*_b_ values in GUVs measured via lsFCS as a function of lipid composition. Each GUV is composed of a particular lipid (i.e., OMPC, POPC, SOPC, DOPC) or a mixture of two lipid species (i.e., SOPC+mSM in 4:1 molar ratio). All GUVs were labelled with 0.005 mol% TF-PC for visualization and lsFCS analysis. **A**: Box plots of the *MMA_b_* values obtained via lsFCS, as a function of lipid composition. **B:** Box plots of the *D_b_* values obtained via lsFCS as a function of lipid composition. For each lipid composition, a total of 53 (up to 122) GUVs were analysed from at least 2 (and up to 5) independent sample preparations (Table S3). The box plots originate from 122, 76, 53, 95 and 72 data points for OMPC, DOPC, POPC, SOPC and SOPC+mSM, respectively (Table S1), within 2 (and up to 5) independent sample preparations. In both panels, each box represents the median (notch), first and third quartile, with ‘x’ marking the mean value. Whiskers indicate standard deviations. ****:p<0.0001 between *D_b_* of DOPC and SOPC+mSM GUVs.

**Figure 2:**
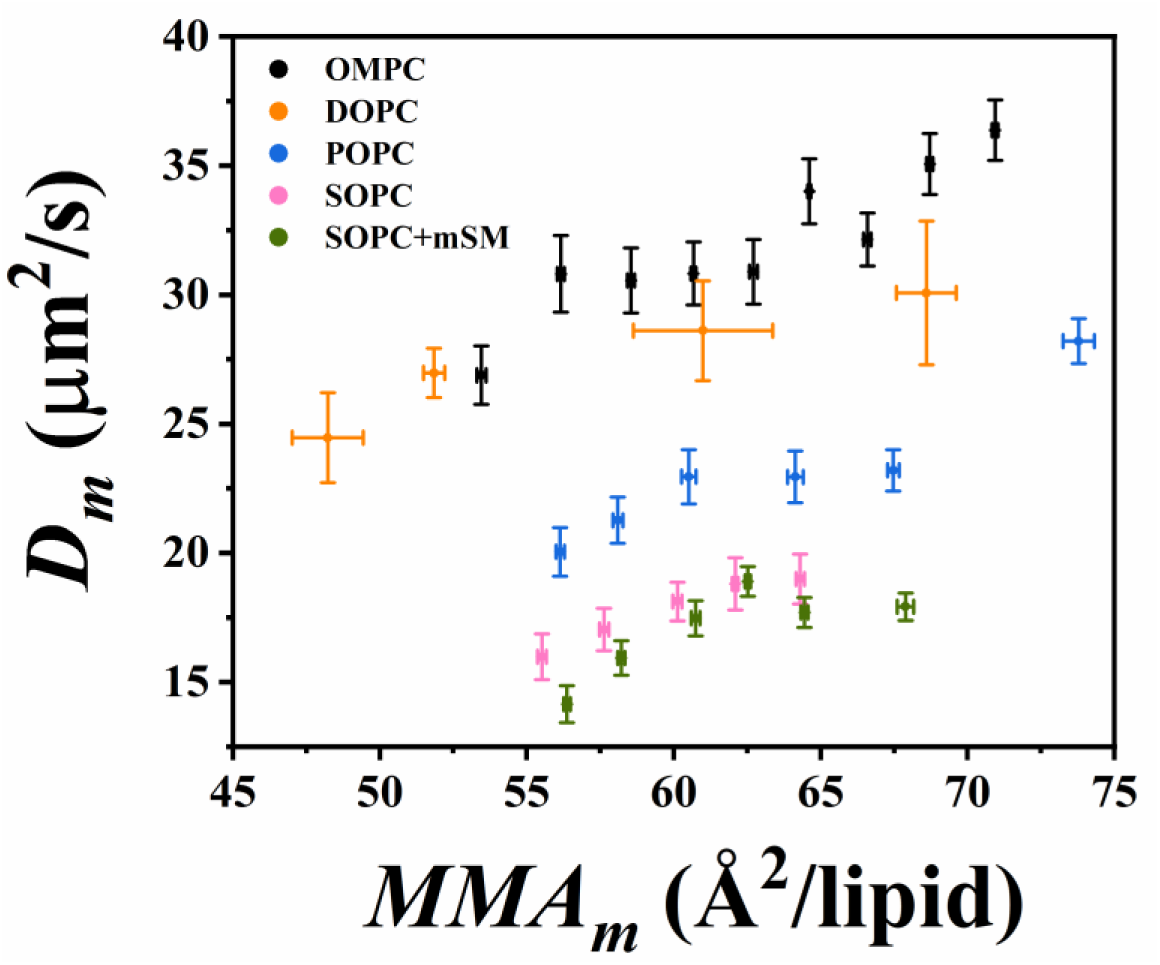
Lipid diffusion coefficients in monolayers formed at air-water interface, measured by RICS. Mean *D_m_* values are shown as a function of *MMA_m_*. Both parameters were obtained via RICS analysis, as described in the main text (section 2.6). Each colour indicates monolayers formed from a specific lipid type, i.e., OMPC, DOPC, POPC, SOPC and SOPC+mSM 4:1 molar ratio. Monolayers were labelled with 0.005 mol% TF-PC. For a better visualization, data points are binned so that each point represents the mean *MMA_m_* and *D_m_* values obtained from 29-32 measurements for OMPC, 3 measurements for DOPC, 16-17 measurements for POPC, 13-22 measurements for SOPC, and 26 measurements for SOPC+mSM. Error bars are standard errors of the mean. For each lipid composition, a total of 9 (up to 19) monolayers were analysed from at least 2 (and up to 5) independent sample preparations. Multiple measurements were performed on each monolayer (Table S2).

**Figure 3:**
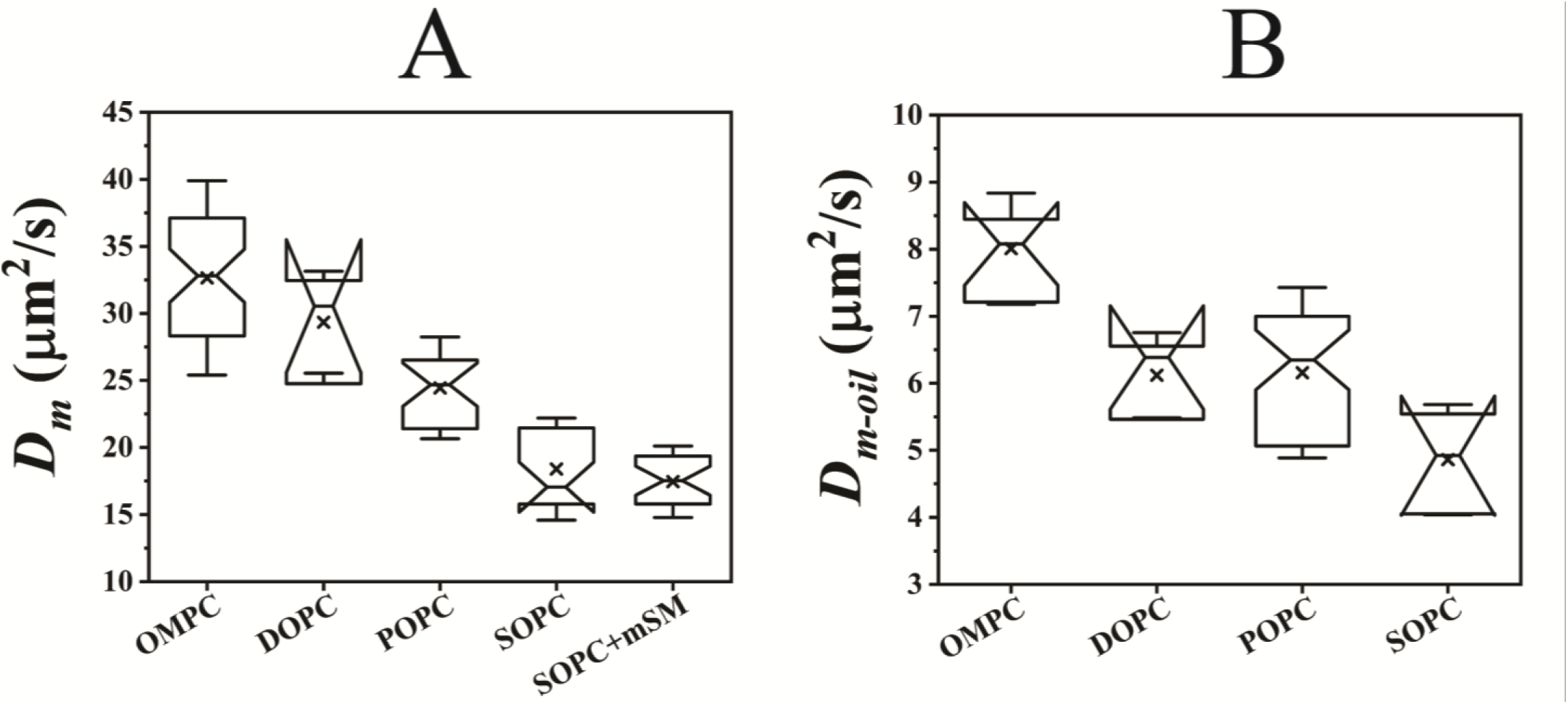
Diffusion coefficients of lipid monolayers measured within specific *MMA_m_* ranges, as a function of lipid composition. **A**: *D_m_* values are obtained via RICS analysis for lipid monolayers composed of OMPC, DOPC, POPC, SOPC and SOPC+mSM (4:1) and labelled with 0.005 mol% TF-PC. All the measured values are shown in Figure 2. In this figure, the box plots represent the spread of *D_m_* values for monolayers with *MMAs* within the range (i.e., one standard error of the mean) of the *MMA_b_* values measured for the corresponding bilayers (Figure 1A). For POPC and DOPC, a larger range is used (i.e., two standard errors of the mean) in order to obtain at least 6 data points for each box. The box plots originate from 49, 6, 25, 23 and 27 data points for OMPC, DOPC, POPC, SOPC and SOPC+mSM, respectively, within 2 (and up to 5) independent sample preparations (Table S2). **B**: *D_m-oil_* values were obtained via lsFCS analysis in pure lipid monolayers of LDs composed of OMPC, DOPC, POPC, SOPC and labelled with 0.001 mol% Rh-PE. Reproducible results and stable samples were obtained only from single lipid mixtures. Like in **A**, the box-and-whisker plots represent the spread of *D_m-oil_* values for monolayers with *MMAs* within the range (i.e. two standard error of the mean) of those measured for the corresponding bilayers (Figure 1A) in order to obtain at least 5 data points for each box. The box plots are composed of 10, 5, 47 and 7 data points for OMPC, DOPC, POPC and SOPC respectively (Table S1). For each lipid composition, a total of 47 (up to 67) LDs were analysed from at least 3 (and up to 5) independent sample preparations (Table S4). In both panels, each box represents the median (notch), first and third quartile with ‘x’ marking the mean value. Whiskers indicate standard deviations.

**Figure 4:**
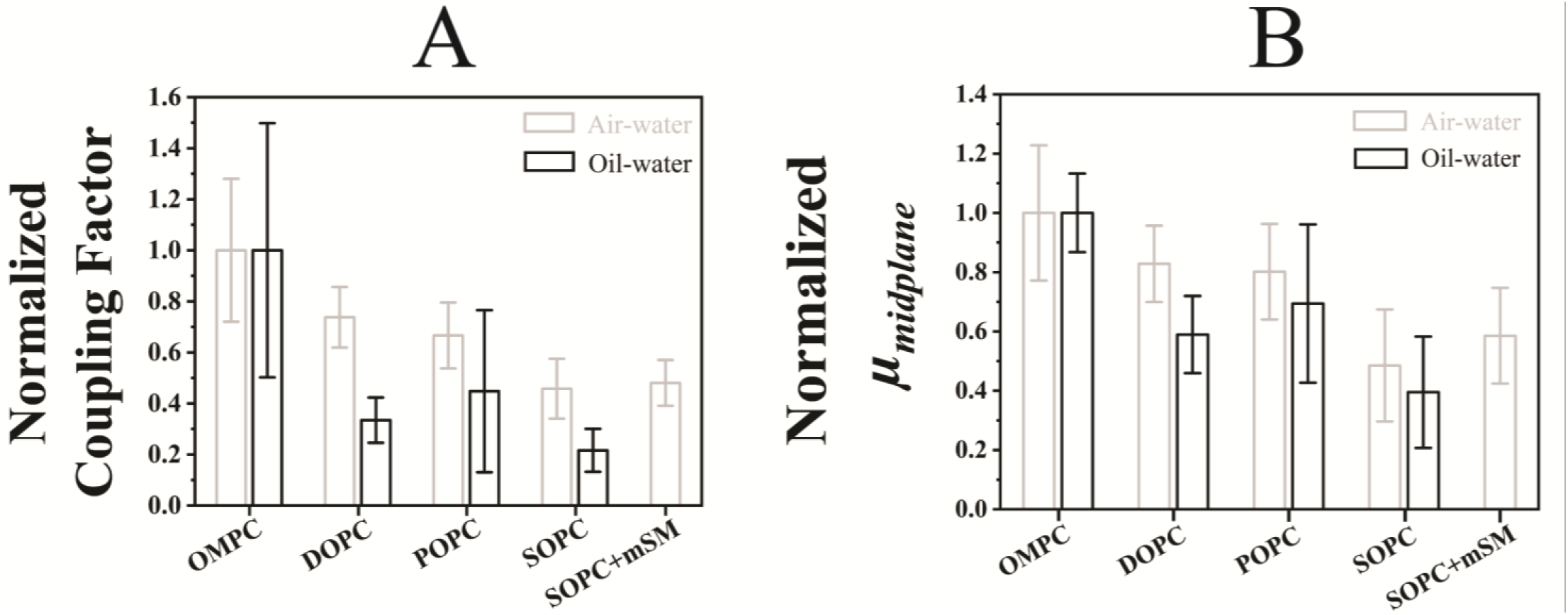
Normalized coupling factors and bilayer midplane viscosities calculated from different monolayer models (air-water and oil-water), as a function of lipid type. **A**: *D_b_* values from Figure 1B, *D_m_* from Figure 3A and *D_m-oil_* from Figure 3B were used to calculate *µ_b_*, *µ_m_* and *µ_m-oil_* using the continuum fluid hydrodynamic model, under the assumption that TF-PC or Rh-PE can be considered a cross-layer particle (see main text, section 2.8). For each lipid composition, the bars represent the mean values of the normalized ratio between the <*µ_b_*> value shown in Table S1 and each *µ_m_* (light grey bars) or *µ_m-oil_* (black bars) value obtained from *D_m_* or *D_m-oil_* values, as represented in Figures 3A and B respectively. **B:** *µ_m_* and *µ_m-oil_* values, and <*D_b_*> from Table S1 were used to calculate the interleaflet/ bilayer midplane viscosities (*µ_midplane_*) with thickness values from the simulation. For each lipid composition, the bars represent the mean values of the normalized *µ_midplane_* obtained from *µ_m_* (light grey bars) and *µ_m-oil_* (black bars). In both panels, the light grey bars originate from 49, 6, 25, 23 and 27 data points for OMPC, DOPC, POPC, SOPC and SOPC+mSM respectively (Table S1); the black bars originate from 10, 5, 47 and 7 data points for OMPC, DOPC, POPC and SOPC respectively (Table S1). The whiskers indicate the standard deviations.

## 3 Results and Discussion

### 3.1 Lipid packing and mobility in GUVs are simultaneously quantified via lsFCS

In order to study the influence of chain length and asymmetry on lipid packing and mobility in bilayers, we prepared GUV model membranes with varying lipid compositions. The lipids for the GUV preparation were chosen based on their similar structure (i.e. one saturated and one mono-unsaturated oleoyl chain) and the difference between the sn-1 and sn-2 acyl chain length (i.e., OMPC, POPC and SOPC). Together with DOPC, these lipids have been previously the subject of investigation in the context of interleaflet interactions ^34,38^. Apart from single-lipid compositions, we examined a lipid mixture as well i.e., GUVs composed of SOPC+mSM, in order to investigate the effect of acyl chain asymmetry ^33,34^.

All GUVs were labelled with 0.005 mol% TF-PC ^94,95^ to enable both membrane visualisation and fluorescence fluctuation measurements ^49^. Specifically, we performed lsFCS to determine the *τ_d_* and *N* of TF-PC molecules within the observation volume, in a single experiment ^49,56,57^. These parameters are used to calculate the diffusion coefficient *D_b_* (equation 2) and *N_total_*. The latter parameter, in turn, allows the estimation of the *MMA_b_* according to equations 3 and 4.

Figures 1 and S2 show that both *MMA_b_* and *D_b_* values do not vary significantly among the tested lipid compositions, taking into account the experimental uncertainties. All average *MMA_b_* values are between ca. 60 and 70 Å^2^/lipid. *D_b_* values are between ca. 6 and 8 µm²/s, with the lowest median values being measured for SOPC and SOPC+mSM and the highest for DOPC (see Table S1).

These results are in good agreement with previously reported data ^30,38,96–101^. It is worth mentioning that the data spread observed in these samples might be partially due to slight variations in membrane tension from vesicle to vesicle ^102,103^, despite the precautions described in section 2.3 employed to obtain GUV samples containing mostly tense/ inflated vesicles.

In conclusion, these experiments indicate that packing density and diffusivity in free-standing planar lipid bilayers can be quantified within a single measurement, using lsFCS.

### 3.2 Diffusive dynamics in monolayers depend on lipid identity and area per lipid

Next, we quantified lipid diffusion in air-water monolayers, i.e. simple models that mimic each leaflet of a bilayer ^104,105^. Previous works have reported *D_m_* values for monolayers with few specific lipid compositions ^43,44,106–108^, however, systematic studies of *D_m_* values as a function of sn-1 and sn-2 acyl chain structure are limited ^108^. Here, monolayers were prepared with the same set of lipids as for GUV samples, using a previously described setup ^42,43^. TF-PC was added in trace amounts (0.005 mol%) for the purpose of visualization and to obtain the parameters *N* and *τ_d_* via fluorescence fluctuation analysis. Using the method described by Khmelinskaia *et al*. ^42^ and Chwastek *et al*. ^43^, the total amount of lipids was adjusted to explore a large range of *MMA* values. The resulting (effective) *MMA_m_* and lipid dynamics were quantified using RICS ^50,66^. Compared to lsFCS, this approach is more suitable for obtaining the required physical parameters for flat lipid systems (e.g. monolayers or bilayers) parallel to the focal plane ^50,67^.

We observed a general positive correlation between *MMA_m_* and *D_m_* (Figure 2), in agreement with published data ^42,43^. Such behaviour is expected, as tighter lipid packing (i.e. lower *MMA_m_*) should decrease lipid diffusive dynamics ^42,43^.

In order to compare lipid dynamics in monolayers and bilayers with the same composition, we focused our analysis on the subset of lipid monolayer preparations which displayed *MMA_m_* values in the range (i.e. within one or two standard errors of the mean) of those measured in average for the corresponding bilayers ^109^ (Figure 1A). Concretely, that implies that all the monolayers examined exhibited similar *MMA_m_* values (i.e., between ca. 60 and 70 Å^2^/lipid). The *D_m_* values obtained for these monolayers are shown in Figure 3A, as a function of lipid composition. For a limited subset of samples (i.e., OMPC and SOPC), we also measured the diffusion of a different fluorophore, i.e. Rh-PE, and found similar *D_m_* values (t-test p=0.56 and 0.73 for *D_m_* of Rh-PE and TF-PC, in OMPC and SOPC monolayers respectively, see Table S1) within similar *MMA_m_* range.

In line with previous results ^92,110^, lipid dynamics are significantly faster in monolayers compared to bilayers by a factor of ca. 2-5. OMPC and DOPC monolayers exhibit the highest *D_m_* values (∼30 µm^2^/s). Reasonably, lipid diffusion is faster in monolayers of fully unsaturated lipids (i.e., DOPC) and slows down for lipids possessing long saturated acyl chains (i.e., *D_m_* for SOPC<POPC<DOPC). Interestingly, within the specific explored *MMA_m_* range, we did not observe significant differences between SOPC monolayers and a more complex mixture including mSM (p=not significant).

Similar experiments were carried out also in LDs, i.e. monolayers at the oil-water interface (Figure 3B and Table S1), as it was previously suggested that such monolayers are more reliable models of the leaflets in a bilayer ^109^. From a qualitative point of view, *D_m-oil_* values vary for the examined lipid compositions following the same trend observed for air-water monolayers (cf. Figures 3A and B). Interestingly, the absolute lipid diffusion values in LDs are ca. 4 times smaller than those measured in air-water monolayers, as expected, due to the high viscosity of the surrounding media (i.e., water and oil vs. water and air) and as it was recently reported for DOPC and POPC oil-water monolayers ^108^.

Finally, as described in section 2.8, *D_m_* and *D_m-oil_* can be used to estimate the monolayer bulk viscosities *µ_m_* and *µ_m-oil_* for these monolayers, according to the continuum fluid hydrodynamic model proposed by Hughes *et al*. ^48,79,85^. As shown in Table S1, the measured viscosities in the monolayer systems are in the range ∼10-60 mPa·s. Furthermore, we do not observe in general significant differences between the viscosity values obtained for each lipid, in air-water monolayers or in oil-water monolayers (see 95% confidence intervals for *µ_m_* and *µ_m-oil_* in Table S1). This result supports the simple view according to which the diffusion of a probe in a lipid monolayer is indeed determined by the bulk viscosity of the monolayer itself (which is within comparable ranges for oil-water and air-water monolayers), the bulk viscosity of the surrounding media and the size of the diffusing molecule ^87^. Previous results have suggested that lipid dynamics in oil-water monolayers might be additionally influenced by interdigitation between oil molecules and lipid acyl chains^111^. Nevertheless, we do not observe such an effect, in the limits of our precision, as evidenced by the similarity of *µ_m_* and *µ_m-oil_* for most lipids (Table S1). Also, it has been suggested that the chemical nature of the diffusing probe (and the specific interactions with the environment) might affect the determination of the membrane viscosity ^87^. Nevertheless, within the experimental uncertainty, we do not observe this effect, as apparent from i) the similar bulk viscosity values obtained in oil-water monolayers and air-water monolayers and ii) the similar *D_m_* values obtained using different fluorophores, at least in OMPC or SOPC monolayers (see above).

### 3.3 Leaflet-leaflet interactions depend on lipid identity

Decreased lipid diffusive dynamics in bilayers, as compared to air-water monolayers with the same *MMA*, can be expected in general considering the different thickness and boundary conditions (water-water for bilayers and, e.g., water-air for monolayers) ^48,79^. Any additional effect might be attributed to leaflet-leaflet interactions. From a qualitative comparison of the results shown in Figures 1B and 3A, it appears that the dynamics of certain lipid species (e.g., OMPC) decrease drastically when switching from an air-water monolayer to a bilayer system. For other lipids (e.g., SOPC), this effect is much smaller. Of note, the large difference between *D_m_* and *D_b_* observed for instance for OMPC samples does not appear to strongly depend on the specific choice of *MMA* (see Figure 2). Such a simple comparison, though, does not take into account other important variables, such as the varying thickness of monolayers and bilayers of different lipid compositions. In what follows, we try thus to quantify such an effect considering also the variations in lipid layer thickness and viscosity of surrounding media, while comparing bilayers and different types of monolayers. The goal of this analysis is not to determine absolute viscosity values for the different analysed lipid systems, but rather to quantify the changes in the “apparent” viscosity and diffusivity in monolayers and bilayers, for different lipid structures.

To this aim, as described in detail in section 2.8, we first calculated the bulk viscosity of the investigated bilayers *µ_b_*, using a simplified approach ^87^, according to which a fluorescent lipid analogue can be approximatively considered a cross-layer particle (i.e., spanning the whole bilayer length, for the purpose of this calculation) ^28,29,33,90–93^. In this case, the viscosities of the surrounding fluids are taken to be that of water on one side and 50 mM sucrose on the other side of the bilayer (see equation 10). As shown in Table S1, the resulting bilayer viscosity values *µ_b_* are between ∼50 and 60 mPa·s, in agreement with previous results^30,112,113^.

Second, we defined a “coupling factor” as the ratio between the mean bilayer bulk viscosity and the bulk viscosity of each corresponding monolayer, for all lipid compositions, as discussed in detail in section 2.8. Figure S3 (A and B) reports the absolute values of the coupling factors calculated for all lipid samples using bulk monolayer viscosities of either air-water or oil-water monolayers. These values range between ca. 1 and 10, with values close to unity (as, e.g., observed for SOPC) suggesting that the bulk viscosity of a bilayer (or each leaflet of the bilayer) is not too far from its corresponding monolayer model. In other words, a value close to 1 would indicate that the decrease in lipid diffusion observed when switching from monolayers to bilayers can simply be ascribed to an increased layer thickness and a change in the surrounding medium, with no additional contribution from e.g. inter-leaflet interactions. Larger values suggest the presence of “additional” factors that might slow down dynamics in bilayers. In order to simultaneously compare the coupling factors referring to air-water and oil-water monolayers, Figure 4A shows the normalized coupling factors obtained from both types of monolayer systems. For the case of SOPC/ mSM mixtures, only water-air monolayers could be reproducibly obtained. In general, no major differences can be observed in the trend of the results obtained from calculations involving either type of monolayer. As previously suggested ^109^ though, we propose that coupling factors calculated using oil-water interfaces might be more reliable than those calculated using air-water monolayers. Regardless, our results suggest that the “additional” decrease in bilayer diffusive dynamics follows the trend OMPC>DOPC≈POPC≳SOPC≈SOPC+mSM.

As an alternative characterization of possible interleaflet interactions causing hindered dynamics in lipid bilayers compared to monolayers, we applied the analysis proposed by Vaz *et al*. ^80^, according to which fluorescent lipids in a bilayer diffuse within a single leaflet. As discussed in detail in section 2.8 (see equation 9), when analysing a single leaflet of a bilayer according to the SPHPW model, the bulk viscosities of the surrounding fluids are that of water on one side of the leaflet and that of the bilayer midplane *µ_midplane_*, on the other. Such “interleaflet or midplane viscosity” is shown in Figure S4 for all the examined lipids, referring alternatively to either *µ_m_* or *µ_m-oil_*. Interestingly, the results obtained for this approach display a trend which is qualitatively similar to that indicated by the “coupling factor” analysis (Figures 4A and B). Independently from the type of monolayer used in the analysis, *µ_midplane_* is consistently the highest in OMPC bilayers. This is particularly evident in the case of oil-water monolayers (Figure S4B). Only minor differences are observed between the other lipid compositions. For example, the addition of mSM to SOPC bilayers does not appear to significantly influence the *µ_midplane_* values (p= not significant). Since mSM is a mixture characterized by a high degree of acyl chain asymmetry ^34,99^, this suggests that interdigitation might not play a major role in explaining specifically the differences in diffusive dynamics between monolayers and bilayers. Of interest, we have previously reported that interdigitation might instead play a role in the process by which reduced dynamics in one leaflet of an asymmetric bilayer induce a similar reduction also in the opposing leaflet ^34^. These observations confirm that acyl chain interdigitation has a non-trivial role in mediating interleaflet interactions and lipid dynamics ^10,39,40^.

In line with the results shown here, previous studies from ours and another group ^34,38^ identified OMPC as a peculiar lipid, in the context of trans-bilayer interactions and spatial organization of the bilayer midplane. The general importance of the spatial distribution of methyl groups at the bilayer midplane in determining interleaflet interaction was also highlighted by other studies ^30,31,40,113^. Specifically, Capponi *et al*. ^38^ proposed that OMPC bilayers are characterized by a “distributed complementarity” of methyl groups, i.e. the peak positions of the *sn*-1 and *sn*-2 methyl distributions in the same leaflet are extraordinarily distant, compared to other lipid bilayers. Of interest, the same behaviour could be reproduced in our simulations that included a ca. 14-fold higher number of lipids in the bilayer (Figure S5). Such spatial distribution of the acyl chain terminal groups might be associated with enhanced packing and, presumably, stronger inter-leaflet interactions across the bilayer midplane. It is worth noting that such behaviour was observed, although to a lower degree, also for SOPC bilayers ^38^. The discrepancy between the results presented in this work for SOPC and those from previous MD investigations should be an object of future investigation.

## 4 Conclusions

In order to verify whether lipid acyl chain structure might influence inter-leaflet interactions, we have systematically compared lipid dynamics in monolayers and bilayers with similar *MMAs*. The expected reduction in diffusive dynamics observed for lipid bilayers appears indeed to depend on the component lipid identity. More in detail, we consistently observed the strongest reduction in diffusive dynamics for bilayers composed of OMPC. On the other hand, the presence of asymmetric lipids (which are supposed to induce strong chain interdigitation) does not appear to affect the decrease of lipid diffusion in bilayers, as compared to monolayers. While our results did not indicate a univocal pattern in how this effect depends on specific structural features of the lipid molecules, there might be a connection between the peculiar behaviour of OMPC and the complementary spatial distribution of terminal methyl ends at the bilayer midplane observed via MD. Such a possibility should be explored through an analogous systematic comparison of lipids with different acyl chains performed via alternative experimental approaches.

## Supporting information

Raw data for all figures

## Abbreviations

OMPC: 1-oleoyl-2-myristoyl-sn-glycero-3-phosphocholine
DOPC: 1,2-dioleoyl-sn-glycero-3-phosphocholine
POPC: 1-palmitoyl-2-oleoyl-glycero-3-phosphocholine
SOPC: 1-stearoyl-2-oleoyl-sn-glycero-3-phosphocholine
mSM: Milk Sphingomyelin
Rh-PE: Rhodamine PE
TF-PC: TopFluor PC
BSA: Bovine serum albumin
PBS: Phosphate buffer saline
NaP: Sodium phosphate
*D*: Diffusion coefficient
*MMA*: Mean molecular area
lsFCS: line-scan fluorescence correlation spectroscopy
RICS: Raster image correlation spectroscopy
PTFE: Polytetrafluoroethylene
LD: Lipid droplet
MD: Molecular dynamics
SDHPW: Saffman-Delbrück-Hughes-Pailthorpe-White
SEM: Standard error of the mean
SD: Standard deviation
*MMA_expected_*: Expected Mean molecular area
*N_Av_*: Avogadro’s number
*r*: radius of the monolayer chamber
*C*: Concentration of lipid solution in molar units
*V*: Volume
*w_0_*: Excitation beam waist
*τ_D_*: Diffusion time
*A_eff_*: Effective illuminated area
*S*: Structural parameter
*MMA_b_*: Mean molecular area of lipid in the bilayer
*D_b_*: Diffusion coefficient of lipids in the bilayer
*N*: Number of fluorescent particles
*N_total_*: Total number of all lipid particles
*MMA_m-oil_*: Mean molecular area of lipid at oil-water interface
*D_m_*: Diffusion coefficient of lipids in the air-water interface
*D_m-oil_*: Diffusion coefficient of lipids in oil-water interface
*MMA_m_*: Mean molecular area of lipid at the air-water interface
*C_%TF-PC_*: Concentration of TF-PC molecules
S/N: Signal to noise
*a*: Radius of diffusing lipid probe head
*h_m_*: Monolayer membrane thickness
*µ_m_*: Air-water monolayer bulk viscosity
*µ_m-oil_*: Oil-water monolayer bulk viscosity
*µ_w_*: Viscosity of water
*µ_s_*: Viscosity of sucrose
*μ_a_*: Viscosity of air
*η_m_*: Monolayer surface viscosity
*ε*: Reduced radius
*h_b_*: Bilayer membrane thickness
*h_l_*: Thickness of a leaflet in a bilayer membrane
*μ_l_*: Bulk viscosity of a leaflet in a bilayer membrane
*μ_midplane_*: Bilayer midplane viscosity
*μ_b_*: Bilayer bulk viscosity

## 5 Author Contributions

**Titas Mandal**: Conceptualization, Methodology, Validation, Formal analysis, Investigation, Data curation, Writing – Original Draft, Writing – Review & Editing, Visualization. **Nadine Brandt**: Methodology, Validation, Investigation, Writing – Review & Editing. **Carmelo Tempra**: Methodology, Software, Validation, Formal analysis, Investigation, Data curation, Writing – Review & Editing, Visualization. **Matti Javanainen**: Methodology, Software, Validation, Formal analysis, Investigation, Resources, Data curation, Writing – Original Draft, Writing – Review & Editing, Visualization, Supervision, Project administration, Funding acquisition. **Balázs Fábián**: Methodology, Software, Validation, Formal analysis, Investigation, Data curation, Writing – Original Draft, Writing – Review & Editing, Visualization. **Salvatore Chiantia**: Conceptualization, Software, Resources, Writing – Original Draft, Writing – Review & Editing, Supervision, Project administration, Funding acquisition.

## Acknowledgements

T.M would like to thank the Postgraduate Scholarship Committee of the University of Potsdam, Germany for his Doctoral dissertation completion scholarship and funding this project. M.J acknowledges support from the Research Council of Finland (postdoctoral researcher grant 338160) and the CSC–IT Center for Science for computational resources. We thank the members of the group for their valuable comments and discussions.

## References

1. Singer, S. J. & Nicolson, G. L. The Fluid Mosaic Model of the Structure of Cell Membranes. Science (1979) 175, 720–731 (1972).

2. Nagle, J. F. & Tristram-Nagle, S. Structure of lipid bilayers. Biochimica et Biophysica Acta (BBA) -Reviews on Biomembranes 1469, 159–195 (2000).

3. Goñi, F. M. The basic structure and dynamics of cell membranes: An update of the Singer–Nicolson model. Biochimica et Biophysica Acta (BBA) - Biomembranes 1838, 1467–1476 (2014).

4. Cebecauer, M. et al. Membrane Lipid Nanodomains. Chem Rev 118, 11259–11297 (2018).

5. Owen, D. M., Williamson, D. J., Magenau, A. & Gaus, K. Sub-resolution lipid domains exist in the plasma membrane and regulate protein diffusion and distribution. Nat Commun 3, 1256 (2012).

6. Eggeling, C. et al. Direct observation of the nanoscale dynamics of membrane lipids in a living cell. Nature 457, 1159–1162 (2009).

7. Garg, S., Rühe, J., Lüdtke, K., Jordan, R. & Naumann, C. A. Domain Registration in Raft-Mimicking Lipid Mixtures Studied Using Polymer-Tethered Lipid Bilayers. Biophys J 92, 1263–1270 (2007).

8. Mouritsen, O. G. & Bagatolli, L. A. Lipid domains in model membranes: a brief historical perspective. Essays Biochem 57, 1–19 (2015).

9. Dietrich, C. et al. Lipid Rafts Reconstituted in Model Membranes. Biophys J 80, 1417–1428 (2001).

10. Collins, M. D. Interleaflet coupling mechanisms in bilayers of lipids and cholesterol. Biophys J 94, (2008).

11. May, S. Trans-monolayer coupling of fluid domains in lipid bilayers. Soft Matter 5, 3148 (2009).

12. Fujimoto, T. & Parmryd, I. Interleaflet Coupling, Pinning, and Leaflet Asymmetry—Major Players in Plasma Membrane Nanodomain Formation. Front Cell Dev Biol 4, (2017).

13. Sarmento, M. J., Hof, M. & Šachl, R. Interleaflet Coupling of Lipid Nanodomains – Insights From in vitro Systems. Front Cell Dev Biol 8, (2020).

14. Bergan, J., Dyve Lingelem, A. B., Simm, R., Skotland, T. & Sandvig, K. Shiga toxins. Toxicon 60, 1085– 1107 (2012).

15. Wernick, N. L. B., Chinnapen, D. J.-F., Cho, J. A. & Lencer, W. I. Cholera Toxin: An Intracellular Journey into the Cytosol by Way of the Endoplasmic Reticulum. Toxins (Basel) 2, 310–325 (2010).

16. Iwabuchi, K., Nakayama, H., Iwahara, C. & Takamori, K. Significance of glycosphingolipid fatty acid chain length on membrane microdomain-mediated signal transduction. FEBS Lett 584, 1642–1652 (2010).

17. Skotland, T., Kavaliauskiene, S. & Sandvig, K. The role of lipid species in membranes and cancer-related changes. Cancer and Metastasis Reviews 39, 343–360 (2020).

18. Klokk, T. I., Kavaliauskiene, S. & Sandvig, K. Cross-linking of glycosphingolipids at the plasma membrane: consequences for intracellular signaling and traffic. Cellular and Molecular Life Sciences 73, 1301–1316 (2016).

19. Dinic, J., Ashrafzadeh, P. & Parmryd, I. Actin filaments attachment at the plasma membrane in live cells cause the formation of ordered lipid domains. Biochimica et Biophysica Acta (BBA) - Biomembranes 1828, 1102–1111 (2013).

20. Štefl, M. et al. Dynamics and Size of Cross-Linking-Induced Lipid Nanodomains in Model Membranes. Biophys J 102, 2104–2113 (2012).

21. Johannes, L. & Römer, W. Shiga toxins — from cell biology to biomedical applications. Nat Rev Microbiol 8, 105–116 (2010).

22. Wang, J. et al. Cross-Linking of GM1 Ganglioside by Galectin-1 Mediates Regulatory T Cell Activity Involving TRPC5 Channel Activation: Possible Role in Suppressing Experimental Autoimmune Encephalomyelitis. The Journal of Immunology 182, 4036–4045 (2009).

23. Wang, Q. & London, E. Lipid Structure and Composition Control Consequences of Interleaflet Coupling in Asymmetric Vesicles. Biophys J 115, 664–678 (2018).

24. Kiessling, V., Wan, C. & Tamm, L. K. Domain coupling in asymmetric lipid bilayers. Biochimica et Biophysica Acta (BBA) - Biomembranes 1788, 64–71 (2009).

25. Wan, C., Kiessling, V. & Tamm, L. K. Coupling of Cholesterol-Rich Lipid Phases in Asymmetric Bilayers. Biochemistry 47, 2190–2198 (2008).

26. Collins, M. D. & Keller, S. L. Tuning lipid mixtures to induce or suppress domain formation across leaflets of unsupported asymmetric bilayers. Proceedings of the National Academy of Sciences 105, 124–128 (2008).

27. Eicher, B. et al. Intrinsic Curvature-Mediated Transbilayer Coupling in Asymmetric Lipid Vesicles. Biophys J 114, 146–157 (2018).

28. Hill, R. J. & Wang, C. Y. Diffusion in phospholipid bilayer membranes: Dual-leaflet dynamics and the roles of tracer-leaflet and inter-leaflet coupling. *Proceedings of the Royal Society A: Mathematical*, Physical and Engineering Sciences 470, (2014).

29. Goutaland, Q. & Fournier, J.-B. Saffman-Delbrück and beyond: A pointlike approach. The European Physical Journal E 42, 156 (2019).

30. Zgorski, A., Pastor, R. W. & Lyman, E. Surface Shear Viscosity and Interleaflet Friction from Nonequilibrium Simulations of Lipid Bilayers. J Chem Theory Comput 15, 6471–6481 (2019).

31. Anthony, A. A., Sahin, O., Yapici, M. K., Rogers, D. & Honerkamp-Smith, A. R. Systematic measurements of interleaflet friction in supported bilayers. Biophys J 121, 2981–2993 (2022).

32. Schoch, R. L., Barel, I., Brown, F. L. H. & Haran, G. Lipid diffusion in the distal and proximal leaflets of supported lipid bilayer membranes studied by single particle tracking. J Chem Phys 148, (2018).

33. Horner, A., Akimov, S. A. & Pohl, P. Long and Short Lipid Molecules Experience the Same Interleaflet Drag in Lipid Bilayers. Phys Rev Lett 110, 268101 (2013).

34. Chiantia, S. & London, E. Acyl Chain length and saturation modulate interleaflet coupling in asymmetric bilayers: Effects on dynamics and structural order. Biophys J 103, 2311–2319 (2012).

35. Otosu, T. & Yamaguchi, S. Reduction of glass-surface charge density slows the lipid diffusion in the proximal leaflet of a supported lipid bilayer. J Chem Phys 151, (2019).

36. Zhang, L. & Granick, S. Interleaflet Diffusion Coupling When Polymer Adsorbs onto One Sole Leaflet of a Supported Phospholipid Bilayer. Macromolecules 40, 1366–1368 (2007).

37. Zhang, L. & Granick, S. Lipid diffusion compared in outer and inner leaflets of planar supported bilayers. J Chem Phys 123, (2005).

38. Capponi, S., Freites, J. A., Tobias, D. J. & White, S. H. Interleaflet mixing and coupling in liquid-disordered phospholipid bilayers. Biochim Biophys Acta Biomembr 1858, 354–362 (2016).

39. Schram, V. & Thompson, T. E. Interdigitation does not affect translational diffusion of lipids in liquid crystalline bilayers. Biophys J 69, 2517–2520 (1995).

40. Den Otter, W. K. & Shkulipa, S. A. Intermonolayer friction and surface shear viscosity of lipid bilayer membranes. Biophys J 93, 423–433 (2007).

41. Zhang, S. & Lin, X. Lipid Acyl Chain cis Double Bond Position Modulates Membrane Domain Registration/Anti-Registration. J Am Chem Soc 141, (2019).

42. Khmelinskaia, A., Mücksch, J., Conci, F., Chwastek, G. & Schwille, P. FCS Analysis of Protein Mobility on Lipid Monolayers. Biophys J 114, 2444–2454 (2018).

43. Chwastek, G. & Schwille, P. A monolayer assay tailored to investigate lipid-protein systems. ChemPhysChem 14, 1877–1881 (2013).

44. Forstner, M. B., Käs, J. & Martin, D. Single Lipid Diffusion in Langmuir Monolayers. Langmuir 17, 567–570 (2001).

45. Gudmand, M., Fidorra, M., Bjørnholm, T. & Heimburg, T. Diffusion and Partitioning of Fluorescent Lipid Probes in Phospholipid Monolayers. Biophys J 96, 4598–4609 (2009).

46. Lira, R. B., Steinkühler, J., Knorr, R. L., Dimova, R. & Riske, K. A. Posing for a picture: Vesicle immobilization in agarose gel. Sci Rep 6, (2016).

47. Przybylo, M. et al. Lipid Diffusion in Giant Unilamellar Vesicles Is More than 2 Times Faster than in Supported Phospholipid Bilayers under Identical Conditions. Langmuir 22, 9096–9099 (2006).

48. Saffman, P. G. & Delbrück, M. Brownian motion in biological membranes. Proceedings of the National Academy of Sciences 72, 3111–3113 (1975).

49. Ries, J., Chiantia, S. & Schwille, P. Accurate Determination of Membrane Dynamics with Line-Scan FCS. Biophys J 96, 1999 (2009).

50. Gielen, E. et al. Measuring diffusion of lipid-like probes in artificial and natural membranes by raster image correlation spectroscopy (rics): Use of a commercial laser-Scanning microscope with analog detection. Langmuir 25, 5209–5218 (2009).

51. Gandhi, S. A., Sanders, M. A., Granneman, J. G. & Kelly, C. V. Four-color fluorescence cross-correlation spectroscopy with one laser and one camera. Biomed Opt Express 14, 3812 (2023).

52. Angelova, M. I. & Dimitrov, D. S. Liposome Electro Formation. Faraday Discuss. Chem. SOC vol. 81 (1986).

53. Stein, H., Spindler, S., Bonakdar, N., Wang, C. & Sandoghdar, V. Production of Isolated Giant Unilamellar Vesicles under High Salt Concentrations. Front Physiol 8, (2017).

54. Omrane, M. et al. LC3B is lipidated to large lipid droplets during prolonged starvation for noncanonical autophagy. Dev Cell 58, 1266–1281.e7 (2023).

55. Gandhi, S. A. et al. Methods for making and observing model lipid droplets. Cell Reports Methods 4, 100774 (2024).

56. Tzoneva, R. et al. Effect of erufosine on membrane lipid order in breast cancer cell models. Biomolecules 10, (2020).

57. Ruan, Q., Cheng, M. A., Levi, M., Gratton, E. & Mantulin, W. W. Spatial-temporal studies of membrane dynamics: Scanning fluorescence correlation spectroscopy (SFCS). Biophys J 87, 1260– 1267 (2004).

58. Petrich, A., Dunsing, V., Bobone, S. & Chiantia, S. Influenza A M2 recruits M1 to the plasma membrane: A fluorescence fluctuation microscopy study. Biophys J 120, 5478–5490 (2021).

59. Petrášek, Z. & Schwille, P. Precise measurement of diffusion coefficients using scanning fluorescence correlation spectroscopy. Biophys J 94, 1437–1448 (2008).

60. Daniels, C. R. et al. Fluorescence correlation spectroscopy study of protein transport and dynamic interactions with clustered-charge peptide adsorbents. Journal of Molecular Recognition 25, 435– 442 (2012).

61. Chenyakin, Y. et al. Diffusion coefficients of organic molecules in sucrose–water solutions and comparison with Stokes–Einstein predictions. Atmos Chem Phys 17, 2423–2435 (2017).

62. The Stokes-Einstein law for diffusion in solution. Proceedings of the Royal Society of London. Series A, Containing Papers of a Mathematical and Physical Character 106, 724–749 (1924).

63. Ries, J. & Schwille, P. Fluorescence correlation spectroscopy. BioEssays 34, 361–368 (2012).

64. Petrich, A., Dunsing, V., Bobone, S. & Chiantia, S. Influenza A M2 recruits M1 to the plasma membrane: A fluorescence fluctuation microscopy study. Biophys J 120, 5478–5490 (2021).

65. Yu, L. et al. A Comprehensive Review of Fluorescence Correlation Spectroscopy. Front Phys 9, (2021).

66. Rossow, M. J., Sasaki, J. M., Digman, M. A. & Gratton, E. Raster image correlation spectroscopy in live cells. Nat Protoc 5, 1761–1774 (2010).

67. Auerswald, J. et al. Measuring protein insertion areas in lipid monolayers by fluorescence correlation spectroscopy. Biophys J 120, 1333–1342 (2021).

68. Iriarte-Alonso, M. A., Bittner, A. M. & Chiantia, S. Influenza A virus hemagglutinin prevents extensive membrane damage upon dehydration. BBA Advances 2, (2022).

69. Lee, J. et al. CHARMM-GUI Input Generator for NAMD, GROMACS, AMBER, OpenMM, and CHARMM/OpenMM Simulations Using the CHARMM36 Additive Force Field. J Chem Theory Comput 12, 405–413 (2016).

70. Klauda, J. B. et al. Update of the CHARMM All-Atom Additive Force Field for Lipids: Validation on Six Lipid Types. J Phys Chem B 114, 7830–7843 (2010).

71. Jorgensen, W. L., Chandrasekhar, J., Madura, J. D., Impey, R. W. & Klein, M. L. Comparison of simple potential functions for simulating liquid water. J Chem Phys 79, 926–935 (1983).

72. Durell, S. R., Brooks, B. R. & Ben-Naim, A. Solvent-Induced Forces between Two Hydrophilic Groups. J Phys Chem 98, 2198–2202 (1994).

73. Abraham, M. J. et al. GROMACS: High performance molecular simulations through multi-level parallelism from laptops to supercomputers. SoftwareX 1–2, 19–25 (2015).

74. Essmann, U. et al. A smooth particle mesh Ewald method. J Chem Phys 103, 8577–8593 (1995).

75. Páll, S. & Hess, B. A flexible algorithm for calculating pair interactions on SIMD architectures. Comput Phys Commun 184, 2641–2650 (2013).

76. Bussi, G., Donadio, D. & Parrinello, M. Canonical sampling through velocity rescaling. J Chem Phys 126, (2007).

77. Parrinello, M. & Rahman, A. Polymorphic transitions in single crystals: A new molecular dynamics method. J Appl Phys 52, 7182–7190 (1981).

78. Hess, B. P-LINCS: A Parallel Linear Constraint Solver for Molecular Simulation. J Chem Theory Comput 4, 116–122 (2008).

79. Hughes, B. D., Pailthorpe A N. B. A. & White, D. L. R. The Translational and Rotational Drag on a Cylinder Moving in a Membrane. J. Fluid Mech (1981).

80. Vaz, W. L. C., Clegg, R. M. & Hallmann, D. Translational diffusion of lipids in liquid crystalline phase phosphatidylcholine multibilayers. A comparison of experiment with theory. Biochemistry 24, 781– 786 (1985).

81. Korson, L., Drost-Hansen, W. & Millero, F. J. Viscosity of water at various temperatures. J Phys Chem 73, 34–39 (1969).

82. Bond, W. N. The viscosity of air. Proceedings of the Physical Society 49, 205–213 (1937).

83. Ando, T. & Skolnick, J. On the Importance of Hydrodynamic Interactions in Lipid Membrane Formation. Biophys J 104, 96–105 (2013).

84. Kučerka, N., Nieh, M. P. & Katsaras, J. Fluid phase lipid areas and bilayer thicknesses of commonly used phosphatidylcholines as a function of temperature. Biochim Biophys Acta Biomembr 1808, 2761–2771 (2011).

85. Petrov, E. P. & Schwille, P. Translational diffusion in lipid membranes beyond the Saffman-Delbrück approximation. Biophys J 94, (2008).

86. Calligaris, S., Mirolo, G., Da Pieve, S., Arrighetti, G. & Nicoli, M. C. Effect of Oil Type on Formation, Structure and Thermal Properties of γ-oryzanol and β-sitosterol-Based Organogels. Food Biophys 9, 69–75 (2014).

87. Adrien, V. et al. How to best estimate the viscosity of lipid bilayers. Biophys Chem 281, 106732 (2022).

88. Jähnig, F. What is the surface tension of a lipid bilayer membrane? Biophys J 71, 1348–1349 (1996).

89. Prince, A. et al. Lipid Specificity of the Fusion of Bacterial Extracellular Vesicles with the Host Membrane. J Phys Chem B 128, 8116–8130 (2024).

90. Galla, H.-J., Hartmann, W., Theilen, U. & Sackmann, E. On Two-Dimensional Passive Random Walk in Lipid Bilayers and Fluid Pathways in Biomembranes. J. Membrane Biol vol. 48 (1979).

91. Lu, D., Vavasour, I. & Morrow, M. R. Smoothed acyl chain orientational order parameter profiles in dimyristoylphosphatidylcholine-distearoylphosphatidylcholine mixtures: a 2H-NMR study. Biophys J 68, 574–583 (1995).

92. Schoch, R. L., Brown, F. L. H. & Haran, G. Correlated diffusion in lipid bilayers. Proceedings of the National Academy of Sciences 118, (2021).

93. Derzko, Z. & Jacobson, K. Comparative lateral diffusion of fluorescent lipid analogs in phospholipid multibilayers. Biochemistry 19, 6050–6057 (1980).

94. Münter, R. et al. Dissociation of fluorescently labeled lipids from liposomes in biological environments challenges the interpretation of uptake studies. Nanoscale 10, 22720–22724 (2018).

95. Shirey, C. M., Ward, K. E. & Stahelin, R. v. Investigation of the biophysical properties of a fluorescently modified ceramide-1-phosphate. Chem Phys Lipids 200, 32 (2016).

96. Filippov, A., Orädd, G. & Lindblom, G. Domain Formation in Model Membranes Studied by Pulsed-Field Gradient-NMR: The Role of Lipid Polyunsaturation. Biophys J 93, 3182–3190 (2007).

97. Filippov, A., Orädd, G. & Lindblom, G. Influence of Cholesterol and Water Content on Phospholipid Lateral Diffusion in Bilayers. Langmuir 19, 6397–6400 (2003).

98. Tristram-Nagle, S., Petrache, H. I. & Nagle, J. F. Structure and Interactions of Fully Hydrated Dioleoylphosphatidylcholine Bilayers. Biophys J 75, 917–925 (1998).

99. Frewein, M. P. K. et al. Interdigitation-Induced Order and Disorder in Asymmetric Membranes. J Membr Biol 255, 407–421 (2022).

100. Alwarawrah, M., Dai, J. & Huang, J. A Molecular View of the Cholesterol Condensing Effect in DOPC Lipid Bilayers. J Phys Chem B 114, 7516–7523 (2010).

101. Koukalová, A. et al. Lipid Driven Nanodomains in Giant Lipid Vesicles are Fluid and Disordered. Sci Rep 7, 5460 (2017).

102. Reddy, A. S., Warshaviak, D. T. & Chachisvilis, M. Effect of membrane tension on the physical properties of DOPC lipid bilayer membrane. Biochim Biophys Acta Biomembr 1818, 2271–2281 (2012).

103. Lindblom, G. & Orädd, G. Lipid lateral diffusion and membrane heterogeneity. Biochimica et Biophysica Acta (BBA) -Biomembranes 1788, 234–244 (2009).

104. Hąc-Wydro, K. & Dynarowicz-Łątka, P. Biomedical applications of the Langmuir monolayer technique. *Annales UMCS*, Chemistry 63, (2010).

105. Stefaniu, C., Brezesinski, G. & Möhwald, H. Langmuir monolayers as models to study processes at membrane surfaces. Adv Colloid Interface Sci 208, 197–213 (2014).

106. Baumgart, T. & Offenhäusser, A. Lateral Diffusion in Substrate-Supported Lipid Monolayers as a Function of Ambient Relative Humidity. Biophys J 83, 1489–1500 (2002).

107. Liu, Y., Zheng, X., Guan, D., Jiang, X. & Hu, G. Heterogeneous Nanostructures Cause Anomalous Diffusion in Lipid Monolayers. ACS Nano 16, 16054–16066 (2022).

108. Asfia, S., Seemann, R. & Fleury, J.-B. Phospholipids diffusion on the surface of model lipid droplets. Biochimica et Biophysica Acta (BBA) - Biomembranes 1865, 184074 (2023).

109. Gruen, D. W. R. & Wolfe, J. Lateral tensions and pressures in membranes and lipid monolayers. Biochimica et Biophysica Acta (BBA) - Biomembranes 688, 572–580 (1982).

110. Babayco, C. B. et al. A comparison of lateral diffusion in supported lipid monolayers and bilayers. Soft Matter 6, 5877–5881 (2010).

111. Bacle, A., Gautier, R., Jackson, C. L., Fuchs, P. F. J. & Vanni, S. Interdigitation between Triglycerides and Lipids Modulates Surface Properties of Lipid Droplets. Biophys J 112, 1417–1430 (2017).

112. Fábián, B., Vattulainen, I. & Javanainen, M. Protein Crowding and Cholesterol Increase Cell Membrane Viscosity in a Temperature Dependent Manner. J Chem Theory Comput 19, 2630–2643 (2023).

113. Merkel, R., Sackmann, E. & Evans, E. Molecular friction and epitactic coupling between monolayers in supported bilayers. Journal de Physique 50, 1535–1555 (1989).

