## Supplementary figures and tables for "A comparison of lipid diffusive dynamics in monolayers and bilayers in the context of interleaflet coupling"

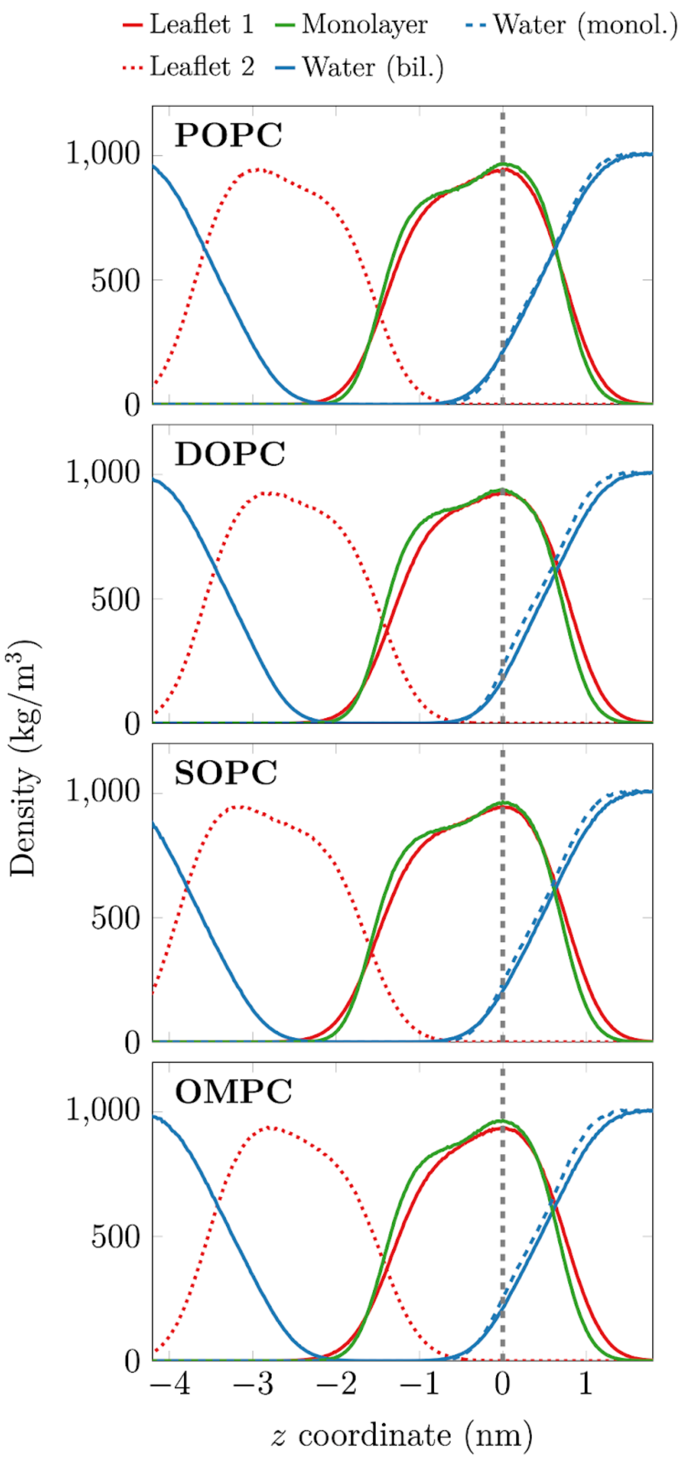


**Figure S1: Density profiles of the simulated lipid bilayers and monolayers.**
The calculation is performed along the *z* coordinate of the simulation box (the direction normal to the lipid layers). For the bilayer, both leaflets are shown separately in solid and dashed lines. The data for the bilayers and monolayers are aligned at the maximal density, and this is shifted to 0. Water density profiles are also shown for both monolayers and bilayers. The visualization allows for the comparison of monolayer and bilayer leaflet thicknesses, the structure of the bilayer and monolayer, and the amount of leaflet interdigitation between the lipid types.

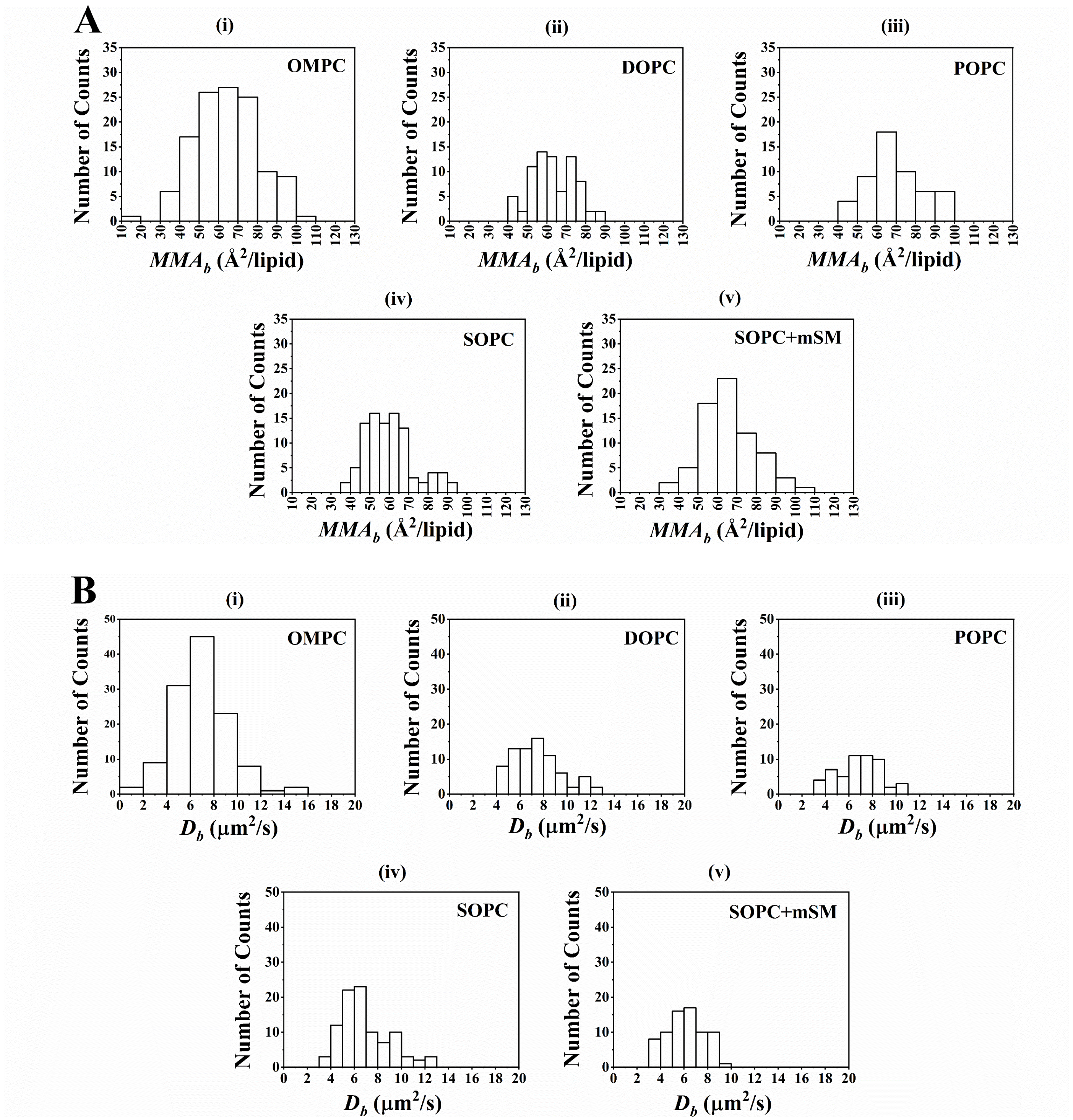


**Figure S2: Histogram distributions of $MMA_b$ and $D_b$ values for GUVs of different lipid compositions.** All measurements of $MMA_b$ and $D_b$ performed in GUVs of different lipid compositions are represented in panels **A** i-v and **B** i-v, respectively. The mean values of these distributions are shown on Table S1. $MMA_b$ (**A**) and and $D_b$ (**B**) values were obtained via lsFCS experiments in GUVs composed of OMPC, DOPC, POPC, SOPC and SOPC+mSM (4:1), labelled with 0.005 mol% TF-PC. The histograms are produced from 122, 76, 53, 95 and 72 data points for OMPC, DOPC, POPC, SOPC and SOPC+mSM respectively (Table S1). For each lipid composition, GUVs were analysed from at least 2 (and up to 5) independent sample preparations (Table S3).

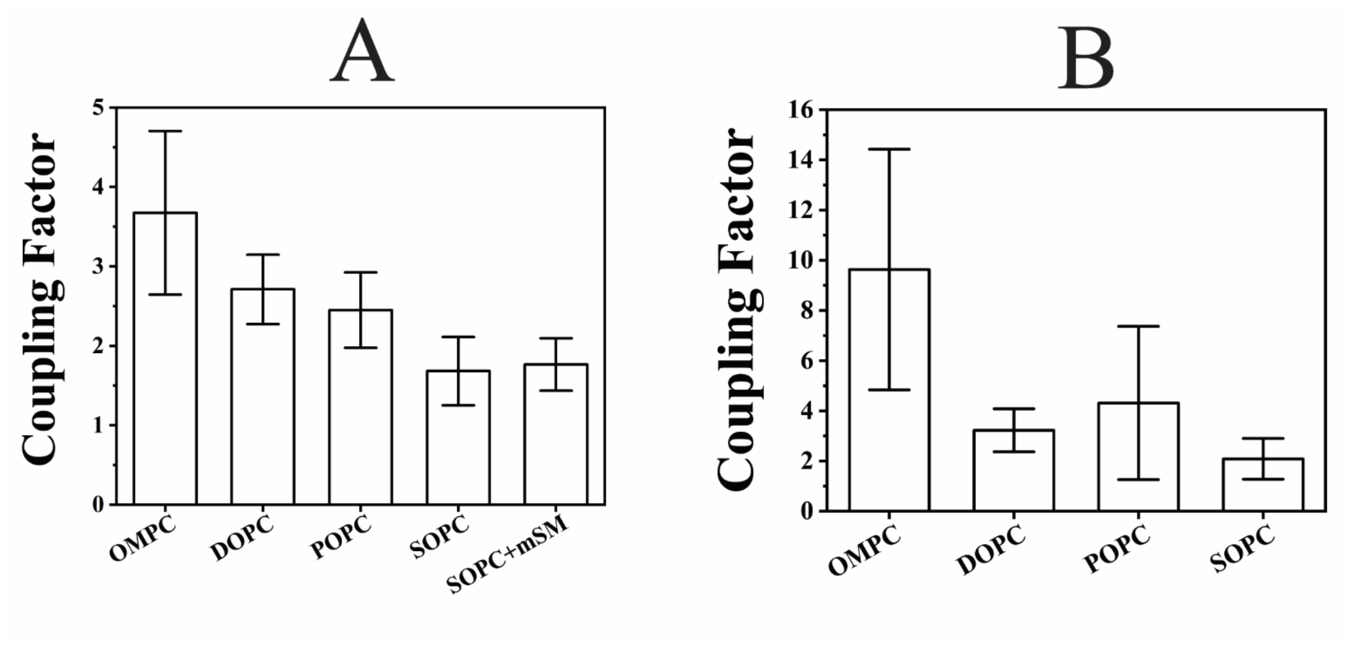


**Figure S3: Non-normalized interleaflet coupling calculated as the ratio of the bulk viscosities of bilayers to that of monolayers.**

$D_b$ values from Figure 1B, $D_m$ from Figure 3A and $D_{m\text{-}oil}$ from Figure 3B were used to calculate the $\mu_b$, $\mu_m$ and $\mu_{m\text{-}oil}$ respectively using the continuum fluid hydrodynamic model, under the assumption that TF-PC or Rh-PE can be considered a cross-layer particle (see main text, section 2.8). For each lipid composition, the bars represent the mean values of the ratio between $<\mu_b>$ value and each $\mu_m$ (**A**) or $\mu_{m\text{-}oil}$ (**B**) obtained from the range of $D_m$ or $D_{m\text{-}oil}$ values, as represented in Figures 3A and B respectively. The bars in **A** are composed of 49, 6, 25, 23 and 27 data points for OMPC, DOPC, POPC, SOPC and SOPC+mSM respectively (Table S1). The bars in **B** are composed of 10, 5, 47 and 7 data points for OMPC, DOPC, POPC and SOPC respectively (Table S1). The whiskers indicate the standard deviations.

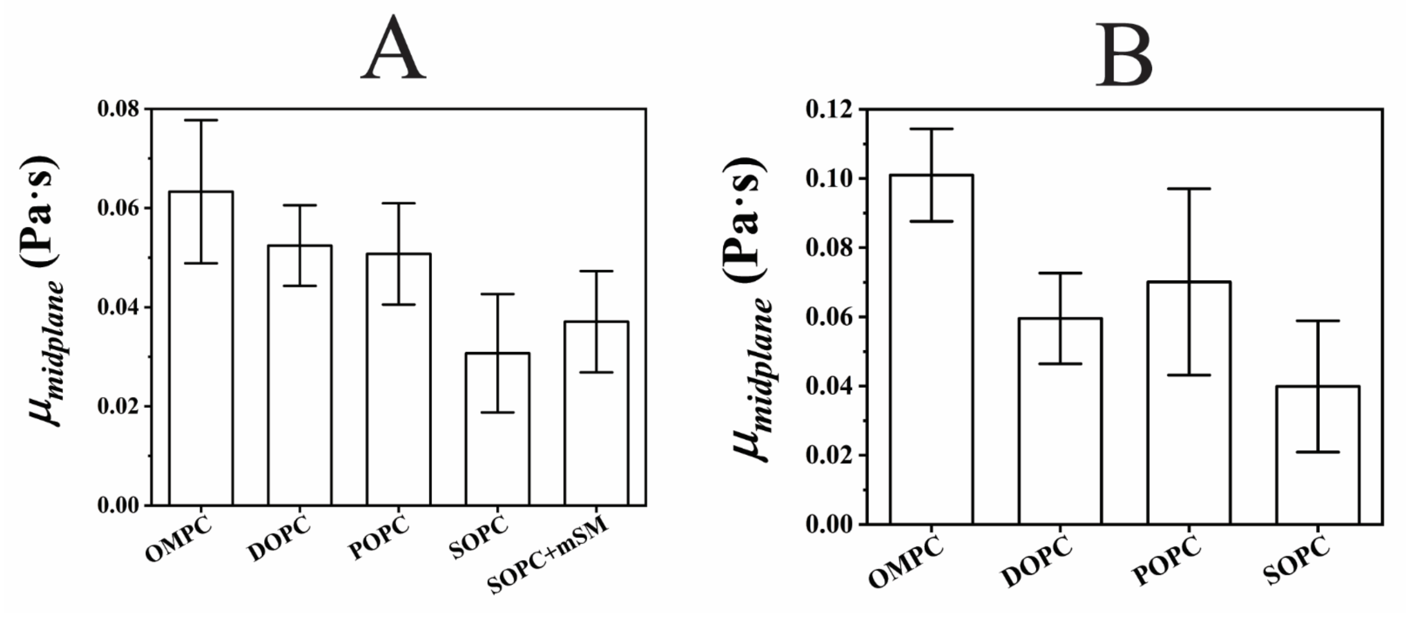


**Figure S4: Bilayer midplane viscosity.**

$\mu_m$ and $\mu_{m\text{-}oil}$ values, and $<D_b>$ from Table S1 were used to calculate the interleaflet/ bilayer midplane viscosities ($\mu_{midplane}$) with thickness values from the simulation. **A** shows the $\mu_{midplane}$ in Pa·s calculated with $\mu_m$ and **B** shows the same calculated with $\mu_{m\text{-}oil}$. In **A**, the bars are composed of 49, 6, 25, 23 and 27 data points for OMPC, DOPC, POPC, SOPC and SOPC+mSM respectively (Table S1). In **B**, the bars are composed of 10, 5, 47 and 7 data points for OMPC, DOPC, POPC and SOPC respectively (Table S1). All the bars represent mean values with whiskers as the standard deviations.

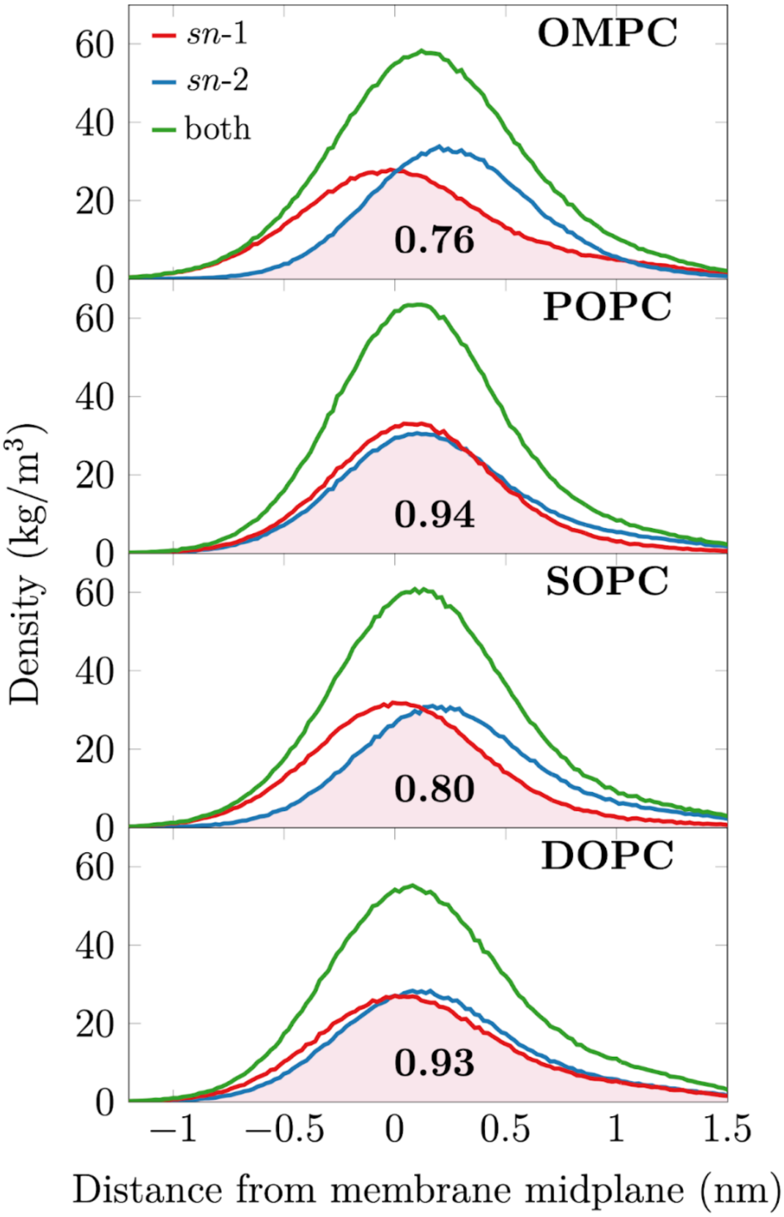


**Figure S5: Density profiles of the terminal carbons of the two acyl chains of a single leaflet shown as a function of distance from the membrane midplane along the *z* coordinate (the direction normal to the membrane plane).**

The profiles indicate the asymmetry of the acyl chains as well as how much each chain is able to penetrate into the opposing leaflet. The membrane midplane is taken as the centre of mass of the entire bilayer. The lipid head group lies in the positive direction (right in the figure). The overlap of the density profiles for the two acyl chains is shaded in pink. The inset numbers indicate the fraction of the total density included in this overlapping part; the larger the number, the more aligned the density profiles for the *sn*-1 and *sn*-2 chains are.

**Table S1:**

| Parameter | OMPC | DOPC | POPC | SOPC | SOPC+mSM |
|---|---|---|---|---|---|
| $MMA_b$ (Å$^2$/lipid)<br><br>Mean±SEM | 64.50±1.50 (n=122)<br><br>SD=16.60 | 63.20±1.30 (n=76)<br><br>SD=11.20 | 69.80±2.0 (n=53)<br><br>SD=14.10 | 60.20±1.30 (n=95)<br><br>SD=12.20 | 65.40±2.0 (n=72)<br><br>SD=13.80 |
| $D_b$ (μm$^2$/s)<br><br>Mean±SEM | 7.10±0.24 (n=122)<br><br>SD=2.70 | 7.46±0.23 (n=76)<br><br>SD=2.0 | 6.80±0.25 (n=53)<br><br>SD=1.80 | 6.86±0.21 (n=95)<br><br>SD=2.10 | 6.15±0.18 (n=72)<br><br>SD=1.50 |
| $D_m$ (μm$^2$/s) Mean±SD | 33±7 (n=49) | 29±4 (n=6) | 24±4 (n=25) | 18±4 (n=23) | 17.40±2.70 (n=27) |
| $D_m$ (μm$^2$/s) Monolayer with Rh-PE Mean±SD | 31±4 (n=7) | | | 17.9±2.4 (n=7) | |
| $D_{m\text{-}oil}$ (μm$^2$/s) Mean±SD | 8.0±0.80 (n=10) | 6.10±0.60 (n=5) | 6.20±1.30 (n=47) | 4.90±0.80 (n=7) | |
| $\mu_m$ (mPa·s) Mean±SD<br><br>95% conf. intervals | 18.20±5.80 (n=49)<br><br>[6.60, 29.80] | 19.60±3.40 (n=6)<br><br>[12.80, 26.40] | 25±5 (n=25)<br><br>[15, 35] | 34.50±8.40 (n=23)<br><br>[17.70, 51.30] | **36±7 (n=27)<br><br>[22, 50] |
| $\mu_{m\text{-}oil}$ (mPa·s) Mean±SD<br><br>95% conf. intervals | 7.50±2.80 (n=10)<br><br>[1.90, 13.10] | 18±5 (n=5)<br><br>[8, 28] | 19±10 (n=47)<br><br>[0, 39] | 30±10 (n=7)<br><br>[10, 50] | |
| $\mu_b$ (mPa·s) Mean±SD<br><br>95% conf. intervals | 62±35 (n=122)<br><br>[0, 132] | 52±17 (n=76)<br><br>[18, 86] | 58±23 (n=53)<br><br>[12, 104] | 55±19 (n=95)<br><br>[17, 93] | **62±21 (n=72)<br><br>[20, 104] |
| Coupling Factor ($<\mu_b>/\mu_m$)<br><br>Mean±SD | 3.70±1.0 (n=49) | 2.70±0.40 (n=6) | 2.50±0.50 (n=25) | 1.70±0.40 (n=23) | 1.80±0.30 (n=27) |
| Coupling Factor ($<\mu_b>/\mu_{m\text{-}oil}$)<br><br>Mean±SD | 10±5 (n=10) | 3.20±0.80 (n=5) | 4±3 (n=47) | 2.10±0.80 (n=7) | |
| $\mu_{midplane}$ (mPa·s) Taking $\mu_m=\mu_l$ (Mean±SD) | 63±14 (n=49) | 52±8 (n=6) | 51±10 (n=25) | 31±12 (n=23) | **37±10 (n=27) |

| | | | | | |
|---|---|---|---|---|---|
| 95% conf. intervals | [35,91] | [36,68] | [31,71] | [7,55] | [17,57] |
| $\mu_{midplane}$ (mPa·s) Taking $\mu_{m\text{-}oil}=\mu_l$ Mean±SD | 101±13 (n=10) [75,127] | 60±13 (n=5) [34,86] | 70±27 (n=47) [16,124] | 40±19 (n=7) [2,78] | |
| Thickness of bilayer (nm) | 5.22 | 5.24 | 5.32 | 5.58 | *5.58 |
| Thickness of each leaflet in bilayer (nm) | 3.01 | 3 | 3.01 | 3.17 | *3.17 |
| Thickness of monolayer (nm) | 2.78 | 2.80 | 2.84 | 2.93 | *2.93 |

*Assumed to be same as that of SOPC
**Calculated using the thickness values of SOPC
95% Confidence intervals are calculated simply as intervals between mean-2SD and mean+2SD

**Table S2:**

| **Parameter** | **OMPC** | **DOPC** | **POPC** | **SOPC** | **SOPC+mSM** |
|---|---|---|---|---|---|
| No. of independent monolayer stocks prepared | 5 | 2 | 3 | 5 | 3 |
| No. of monolayers measured and analysed | 18 | 9 | 16 | 19 | 14 |
| No. of measurements and analysis per monolayer | 11 to 31 | 2 to 12 | 4 to 30 | 5 to 30 | 9 to 33 |

**Table S3:**

| **Parameter** | **OMPC** | **DOPC** | **POPC** | **SOPC** | **SOPC+mSM** |
|---|---|---|---|---|---|
| No. of independent bilayer stocks prepared | 5 | 2 | 3 | 5 | 3 |
| No. of GUV preparations | 7 | 6 | 5 | 5 | 3 |
| No. of measurements and analysis per GUV preparation | 11 to 28 | 5 to 21 | 7 to 14 | 6 to 26 | 23 to 25 |

**Table S4:**

| **Parameter** | **OMPC** | **DOPC** | **POPC** | **SOPC** |
|---|---|---|---|---|
| No. of independent monolayer aliquots made | 4 | 5 | 3 | 4 |
| No. of LD preparations | 4 | 5 | 3 | 4 |
| No. of measurements and analysis per LD preparation | 4 to 21 | 5 to 18 | 10 to 21 | 14 to 19 |
| Total number of LDs measured and analysed | 66 | 54 | 47 | 67 |
